# Cytoplasmic processing of human transfer RNAs

**DOI:** 10.1101/2022.04.28.489951

**Authors:** Yasutoshi Akiyama, Shawn M. Lyons, Takaaki Abe, Paul J. Anderson, Pavel Ivanov

## Abstract

Biogenesis of different classes of eukaryotic RNAs proceeds via different pathways that require strict spatiotemporal resolution. The intracellular organization has evolved to provide order for RNA processing to coordinate different maturation steps with specific enzymatic reaction. In higher eukaryotes, processing of transfer RNAs (tRNAs) is postulated to be almost entirely intranuclear, while in lower eukaryotes like yeast, tRNA maturation is both nuclear and cytoplasmic. Here, we show that tRNA processing is largely cytosolic event in human cells. After transcription, unprocessed precursors of tRNAs (pre-tRNAs) bound by La protein are exported from the nucleus into the cytoplasm. Using cell fractionation analysis and pre-tRNA-specific fluorescence in situ hybridization (FISH) protocol, we show tRNA splicing/ligation and end-processing takes place in the cytoplasm where majority of processing enzymes is also located. We propose here a model where processing of intron-less and intron-containing pre-tRNAs is cytoplasmic similarly to the observed in lower eukaryotes, although tRNA splicing precedes 5’- and 3’-end processing in human cells unlike in yeast.

## INTRODUCTION

All eukaryotic primary tRNA transcripts (pre-tRNAs), which are synthesized in the nucleus by RNA Polymerase III, contain 5’-leader and 3’-trailer sequences, while some pre-tRNAs also contain introns. Interestingly, generation of a complete set of tRNAs for their participation in protein synthesis requires intron removal since tRNA genes in eukaryotes encode at least one tRNA family (tRNA^Tyr^) that is exclusively intron-containing (1). In human cells, intron removal from pre-tRNA requires action of the hetero-tetrameric tRNA <u>s</u>plicing <u>e</u>ndo<u>n</u>uclease (TSEN) complex consisting of TSEN4, TSEN15, TSEN34 and TSEN54 subunits (Sen4, Sen15, Sen34 and Sen54 in yeast, respectively) (2). Catalytic subunits of TSEN complex (TSEN2 and TSEN34) cleave intron-containing pre-tRNAs at 5’- and 3’-splice sites (3), and following the intron removal, ligation of tRNA exon halves is catalyzed by tRNA ligase(s) (4). tRNA ligation in human cells appears to be predominantly dependent on the action of RTCB (HSPC117) ligase, which is found in stable complex with other factors such as Archease, DDX1, CGI-99, FAM98B and ASW (5), although an alternative route relying on the action of the RNA kinase CLP1 is also reported (6). Finally, for both intron-less and intron-containing tRNAs, 5’-leader and 3’-trailer sequences are removed post-transcriptionally by ribonuclease P (RNase P) and RNase Z, followed by addition of the universal CCA triplet to the 3’-end of the tRNA transcript (3’-CCA). Numerous nucleotides of pre-tRNA are also modified in the nucleus before primary tRNA export, followed by addition of other modifications in the cytoplasm to complete tRNA maturation (extensively reviewed in (7-10)).

It should be noted that the process of tRNA maturation in lower (e.g., yeast) and higher (e.g., mammals) eukaryotes significantly differs both in the location of processing steps and the order of maturation (reviewed in (7-10)). First, it is postulated that most of the processing steps in vertebrates happen in the nucleus (nucleoplasm), while in yeast some steps of pre-tRNA processing such as pre-tRNA splicing occur in the cytoplasm. Second, it is postulated that tRNA splicing occurs before the end maturation in vertebrates, while in yeast splicing occurs after 5’-leader/3’-trailer removal and the addition of 3’-CCA (reviewed in (11)).

At the same time, our understanding of tRNA biogenesis in human cells is quite incomplete and fragmented. Most of the currently available data on mechanisms of tRNA biogenesis is predominantly based on the studies from lower eukaryotes and earlier studies based on the microinjection of pre-tRNA genetic constructs into the nucleus of Xenopus eggs (12), which are non-dividing, non-human, specialized cells with number of physiological differences to proliferating cells. Moreover, available literature also suggests that some of protein factors implicated in human tRNA biogenesis (e.g., RTCB), can be found both in the nucleus and the cytoplasm (based in immunofluorescence studies) (13,14). This could suggest that some steps of human pre-tRNA splicing may happen in the cytoplasm, or/and these nucleocytoplasmic factors have other functions besides tRNA-related (reviewed in (15)).

Our work here is entirely based on the analysis of endogenous pre-tRNA species and mature tRNAs without the use of genetic constructs. We employed cell fractionation and pre-tRNA FISH protocols to demonstrate that tRNA precursors can predominantly be detected in the cytoplasm where they undergo processing. In addition, the analysis of subcellular distribution of the endogenous factors involved in tRNA maturation (i.e., without overexpression of artificially tagged proteins) also suggests that several enzymes involved in pre-tRNA processing are found in the cytoplasmic compartment. Our data suggest that in human cells, tRNA splicing precedes 5’- and 3’-end maturation unlike in lower eukaryotes. Finally, we show that nucleocytoplasmic protein La contributes to the export of pre-tRNAs into the cytoplasm. Altogether, our work uncovers new aspects of pre-tRNA maturation in human cells.

## MATERIAL AND METHODS

### Cell culture and transfection

U2OS, 293T and RPE-1 cells were purchased from the American Type Culture Collection (ATCC). U2OS and 293T cells were cultured at 37°C in a CO_2_ incubator in Dulbecco’s modified Eagle’s medium (DMEM) supplemented with 10% fetal bovine serum (FBS) (Sigma-Aldrich) and 1% of penicillin/streptomycin (Sigma-Aldrich). RPE-1 cells were maintained in DMEM/F-12 supplemented with 10% FBS and 1% of penicillin/streptomycin. For knockdown experiments, cells were transfected with siRNA at a concentration of 40 nM using Lipofectamine 2000 (Invitrogen) according to the manufacturer’s protocol for reverse transfection, then collected 72 hours after transfection. The siRNAs (siGENOME SMARTpool for RTCB and ON-TARGETplus SMARTpool for CLP1, TSEN2, XPOT, XPO1, NXF1, RAN and negative control) were purchased from GE Dharmacon.

### shRNA-mediated knockdown

The following plasmids were purchased from Addgene (Cambridge, MA, USA): pLKO.1 (#10878), pMD2G (#12259) and psPAX2 (#12260). Oligonucleotides encoding shRNAs were synthesized by IDT (Integrated DNA Technologies, Coralville, IA) and cloned into pLKO.1 vector following manufacturer’s instruction. The lentiviral particles were produced by co-transfection of 293T cells with three plasmids (pLKO.1, pMD2G and psPAX2 vectors) using Lipofectamine 2000 (Invitrogen). Cells were infected with lentiviral particles in the presence of 8 µg/ml polybrene (Sigma-Aldrich) overnight and were selected one day after viral transduction with 1.5 µg/ml of puromycin (Sigma-Aldrich). Cells were collected 7 days after transduction and subjected to subcellular fractionation described below. The sequences of the shRNA-encoding oligonucleotides were shown in Supplementary Table S1.

### Subcellular fractionation

Subcellular fractionation was performed according to the method previously reported (16) with slight modifications. For isolation of fractionated RNAs, cells were resuspended in 380 µl of ice-cold Hypotonic Lysis Buffer (HLB) [10 mM Tris-HCl (pH 7.5), 10 mM NaCl, 3 mM MgCl_2_, 0.3%(vol/vol) NP-40 and 10%(vol/vol) glycerol] supplemented with RNasin (Promega). Ten minutes after incubation on ice, cells were centrifuged and the supernatant (cytoplasmic fraction) was mixed with 650 µl of Trizol-LS (Invitrogen). Pellets (nuclei) were washed with 1 mL of HLB for three times, then 1 mL of Trizol (Invitrogen) was added directly into the pellets and vortexed thoroughly. Isolated RNAs were eluted in the same amount of water.

For isolation of fractionated protein samples, cells were resuspended in 500 µl of ice-cold HLB supplemented with Halt Protease Inhibitor Cocktail (ThermoFisher Scientific) and Halt Phosphatase Inhibitor Cocktail (ThermoFisher Scientific). Ten minutes after incubation on ice, cells were centrifuged and the supernatant (cytoplasmic fraction) was transferred into a new tube, then 5M NaCl was added to give a final concentration of 150 mM. The pellets were washed with 1 mL of HLB for three times, then resuspended in 500 µl of Nuclear Lysis Buffer (NLB) [20 mM Tris-HCl (pH7.5), 150 mM KCl, 3 mM MgCl_2_, 0.3% (vol/vol) NP-40 and 10% (vol/vol) glycerol] supplemented with Halt Protease Inhibitor Cocktail and Phosphatase Inhibitor Cocktail. The suspension was sonicated on ice and then centrifuged. The supernatant (Nuclear fraction) was transferred into a new tube.

### Northern blotting

Total RNA was extracted by using Trizol (Invitrogen). RNA (7.5 μg per well) was run on 10% TBE-urea gels (ThermoFisher Scientific), transferred to positively charged nylon membranes (Roche), and hybridized overnight at 40°C with digoxigenin (DIG)-labeled DNA probes in DIG Easy Hyb solution (Roche). After low stringency washes (washing twice with 2× SSC/0.1% SDS at room temperature) and high stringency wash (washing once with 1× SSC/0.1% SDS at 40°C), the membranes were blocked in blocking reagent (Roche) for 30 min at room temperature, probed with alkaline phosphatase-labeled anti-digoxigenin antibody (Roche) for 30 min, and washed with 1x TBS-T. Signals were visualized with CDP-Star ready-to-use (Roche) and detected using ChemiDoc imaging system (BioRad) according to the manufacturer’s instructions. Densitometry was performed using ImageJ software (NIH). Oligonucleotide probes were synthesized by IDT. DIG-labeled probes were prepared using the DIG Oligonucleotide tailing kit (2nd generation; Roche) according to the manufacturer’s instructions. The sequences of the probes were shown in Supplementary Table S2.

### Western Blotting

Cell lysates were prepared using RIPA buffer containing protease inhibitor cocktail (ThermoFischer Scientific). Protein lysate was run on a 4–20% SDS PAGE and transferred to nitrocellulose membrane (BioRad). Blots were blocked with 5% skim milk in TBS-T (Tris-buffered saline with 0.1% Tween 20) for 1 hr at room temperature, then incubated overnight at 4 °C with one of the antibodies listed in Supplementary Table 2. After washing and incubation for 1 h at room temperature with HRP-conjugated secondary antibody (Jackson ImmunoResearch), the protein signal was detected using a SuperSignal West Pico PLUS Chemiluminescent Substrate (ThermoFisher Scientific) and ChemiDoc imaging system (BioRad) according to the manufacturer’s instructions. A list of the antibodies used in this study was shown in Supplementary Table S3.

### Immunofluorescence

Immnunofluorescence was performed as previously described (17) with slight modifications. Cells were grown on coverslips, washed with PBS, fixed with 4% paraformaldehyde in PBS for 15 min at room temperature, then permeabilized with 0.1% Triton X-100 in PBS for 10 min. Coverslips were blocked with 5% normal horse serum in PBS (NHS-PBS) at room temperature for 30 min, then incubated overnight with primary antibodies diluted at 1:200 with NHS-PBS at 4 °C. After washing, cells were incubated with NHS-PBS containing fluorophore-conjugated secondary antibodies (Jackson ImmunoResearch) and Hoechst dye at room temperature for 1 hr. Cells were washed and mounted with vinol mounting media.

In order to reveal TSEN2 antigens, heat-induced antigen retrieval (HIER) (18) was performed by incubating coverslips in citrate buffer (10mM citric acid, 0.05% Tween 20, pH 6.0) at 95 °C for 10 min before blocking.

### Fluorescence in situ hybridization (FISH) for pre-tRNA

FISH for pre-tRNAs was performed as previously described (17) with slight modifications. Briefly, cells were fixed with 4% paraformaldehyde in PBS for 15 min then permeabilized with 0.1% Triton X-100 in PBS for 10 min. Cells were then incubated overnight in 70% ethanol at 4 °C. The following day, cells were washed twice with 2x saline sodium citrate (SSC) buffer, then pre-hybridized in hybridization buffer (Sigma-Aldrich) at 42 °C for 30 min. After replacing hybridization buffer with 4 ng/μl of biotinylated probes diluted in hybridization buffer, samples were heated at 76 °C for 5 min in humidified chamber to denature target pre-tRNAs, then hybridized at 42 °C for 1 hr in humidified chamber. After extensive washes with 2x SSC buffer at 37 °C, the hybridized probe was detected via fluorophore-conjugated streptavidin (Jackson ImmunoResearch), followed by immunostaining when needed as described above. A list of the biotinylated probes used in this study is shown in Supplementary Table S4.

### Ribonucleoprotein immunoprecipitaion and Co-immunoprecipitation using anti-La antibody

The isolation of cytoplasmic La-associated RNA was performed according to the protocol described by Keene et al (19). Briefly, U2OS cells were suspended and incubated in Polysome Lysis Buffer (PLB) [100 mM KCl, 5 mM MgCl_2_, 10 mM HEPES (pH 7.0), 0.5% NP-40, 1 mM DTT and 400 µM vanadyl ribonucleoside complexes (VRC; NEB)] supplemented with RNasin and Halt Protease Inhibitor Cocktail on ice for 10 min. Then the lysate was frozen until use. The thawed lysate was centrifuged to pellet nuclei. The supernatant (cytoplasmic lysate) was diluted 10 times with NT2 buffer [50 mM Tris-HCl (pH 7.4), 150 mM NaCl, 1 mM MgCl_2_, 0.05% NP-40] supplemented with RNasin, and incubated with mouse anti-La antibody (312B, Santa Cruz) or mouse IgG-1κ (for negative control) conjugated to Protein A/G UltraLink Resin (Thermo Fisher Scientific) at room temperature for 2 h with gentle rotation. After washing the beads three times with NT2 buffer, RNA was extracted from the beads using Trizol. The input cytoplasmic fraction and the supernatant fraction were also subjected to RNA isolation using Trizol.

Alternatively, proteins were eluted from the beads by adding SDS sample buffer followed by boiling. The input cytoplasmic fraction and the supernatant fraction were also collected and subjected to Western blotting.

## RESULTS

### Precursor tRNAs are enriched in the cytoplasm

Nuclear/cytoplasmic fractionation of human osteosarcoma U2OS cells (Figure 1A) reveals strong enrichment of mature tRNAs in the cytoplasmic fraction. The specificity of fractionation has been validated by northern blot (NB) analysis of RNA species, which are exclusively nuclear (snRNAs, Figure 1B) or cytoplasmic (Y RNAs, Figure 1B). Analysis of nuclear and cytoplasmic fractions using NB against both intron-less (tRNA^iMet-CAT^ and tRNA^Gly-GCC^) and intron-containing (tRNA^Tyr-GTA^ and tRNA^Leu-CAA^) tRNAs shows that both subclasses of pre-tRNAs are predominantly present in the cytoplasmic fraction (Figure 1C). Pre-tRNA FISH analysis using probes against different parts of pre-tRNA (e.g., intron of tRNA^Leu-CAA^, or 3’-trailers of tRNA^Tyr-GTA^ and tRNA^Gly-GCC^, Figure 1D) confirmed cytoplasmic localization of various pre-tRNA in agreement with subcellular fractionation studies. We further included other cell lines, e.g. human embryonic kidney HEK293 and human retinal pigment epithelial RPE-1 cells in our analysis, and confirmed predominant cytoplasmic localization of pre-tRNAs (Supplementary Figure S1). Specificities of probes used for pre-tRNA FISH were also validated by NB, their sensitivity to the RNase A treatment and denaturation by temperature (Supplementary Figure S2A-C). We also used RNA Pol III inhibitor (Cas 577784-91-9) to determine how inhibition of tRNA genes transcription affects detection of pre-tRNAs by NB and tRNA FISH. Pol III inhibitor does not change signal for mature tRNAs suggesting that mature tRNA levels are not affected (Supplementary Figure S2D). In contrast, levels of pre-tRNAs of both intron-containing tRNA^Tyr-GTA^ and intron-less tRNA^Gly-GCC^ are down-regulated suggesting that (1) the inhibitor effectively shuts off transcription of tRNA genes, and (2) that our NB probes are specific to pre-tRNAs. These results were further confirmed by the analysis using pre-tRNA FISH in the presence or absence of the Pol III inhibitor (Supplementary Figure S2D, E).

**Figure 1.**
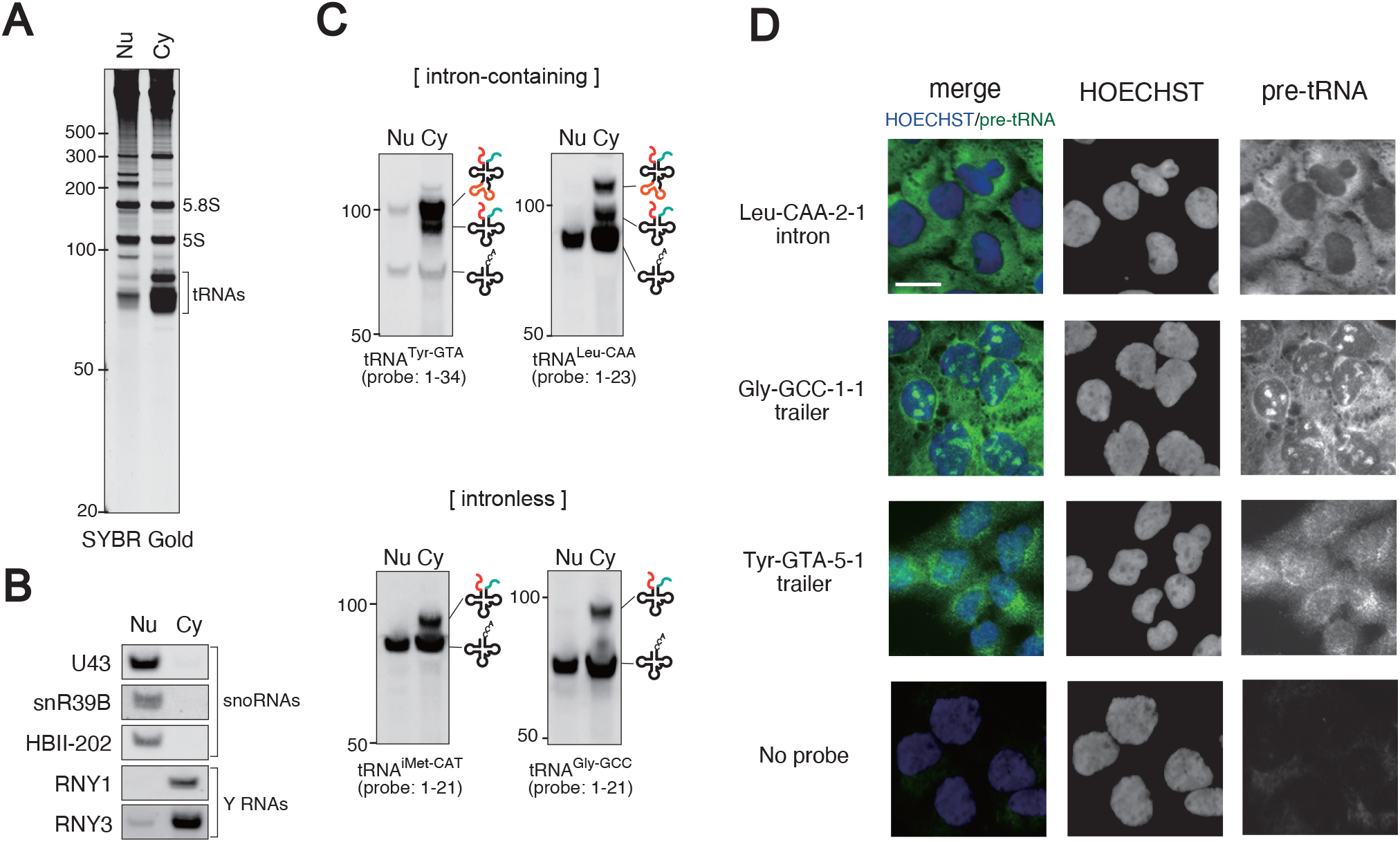
Precursor tRNAs are localized in the cytoplasm. (A-C) Enrichment of pre-tRNAs in the cytoplasmic fraction. (A) SYBR Gold staining of fractionated RNAs. (B) Validation of fractionation method. Nuclear RNAs (snoRNAs) and Cytoplasmic RNAs (Y-RNAs) are enriched in the nuclear and cytoplasmic fraction, respectively. (C) pre-tRNAs are enriched in the cytoplasmic fraction regardless of whether they are intron-containing or intron-less genes. (D) Fluorescent in situ hybridization (FISH) confirming cytoplasmic localization of pre-tRNAs.

### Components of pre-tRNA processing machinery are also enriched in the cytoplasm

We further extended our subcellular fractionation and immunofluorescence (IF) studies to determine whether some components of tRNA processing machinery are also cytoplasmic. Specifically, we analyzed components of tRNA splicing endonuclease TSEN complex (20), the RNA kinase CLP1 (21), RTCB ligase and its co-factors DDX1 and Archease (ARCH) (4), subunits of the RNase P and its RNA component H1 RNA (22), RNase Z1 (23), and CCA-adding enzyme TRNT1 (24). As controls for the fractionation, we used Lamin B1 and alpha-tubulin as nuclear and cytoplasmic markers, respectively (Figure 2A). Most of the analyzed protein factors are found in both nuclear and cytoplasmic fractions, although some components of pre-tRNA processing machinery, e.g. TSEN complex subunits, are enriched in the cytoplasmic fraction (Figure 2A and Supplementary Figure S3). In addition to subcellular fractionation studies, we determined subcellular localization of these factors by IF. For example, and in agreement with Figure 2A, TSEN2 and TRNT1 show both nuclear and cytoplasmic IF staining (Figure 2B-E, Supplementary Figure S4). Importantly, their knockdown using siRNAs (Figure 2B and 2D, respectively), decreases their cytoplasmic (and not nuclear) signal (Figure 2C and 2E, respectively). In addition, we detected H1 RNA, the RNA component of human RNase P, in the cytoplasm using three NB probes for different parts of the molecule (Figure 2F) and by FISH (Figure 2G).

**Figure 2.**
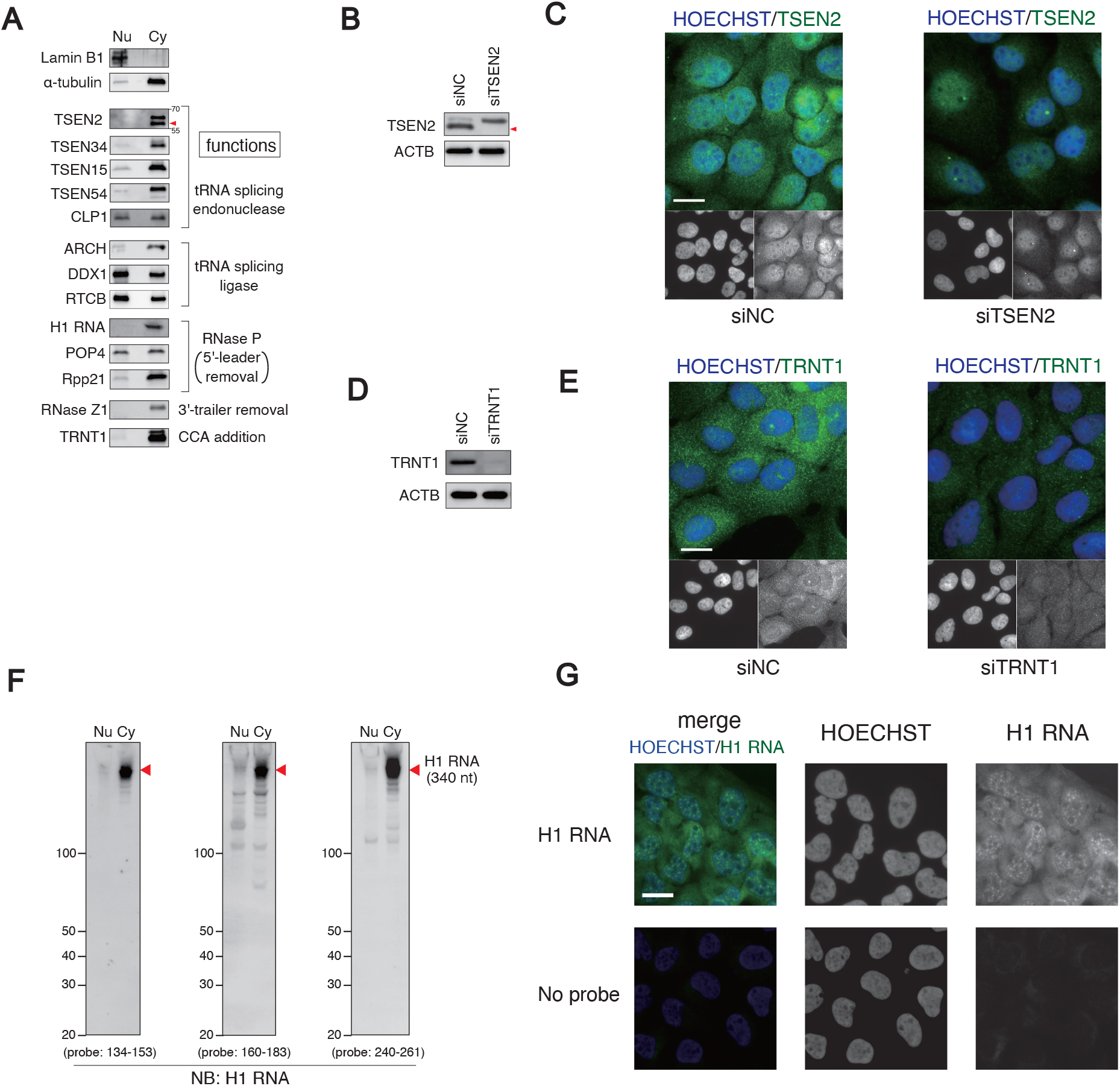
Components of pre-tRNA processing are also enriched in the cytoplasm. (A) Subcellular fractionation confirming enrichment of pre-tRNA processing components in the cytoplasmic fraction. (B-C) TSEN2 subunit localizes in the cytoplasm. (B) siRNA-mediated knockdown of TSEN2 subunit. (C) Cytoplasmic TSEN2 signal is decreased by TSEN2 knockdown. (D-E) CCA-adding enzyme TRNT1 also localizes in the cytoplasm. (D) siRNA-mediated knockdown of TRNT1. (E) Cytoplasmic TRNT1 signal is decreased by TRNT1 knockdown. (F-G) Cytoplasmic localization of H1 RNA. (F) Subcellular fractionation combined with Northern blotting for H1 RNA. All the three probes showed the cytoplasmic enrichment. (G) FISH for H1 RNA shows the cytoplasmic localization. The mixture of three probes used in (F) was used as FISH probes.

### tRNA splicing takes place in the cytoplasm and precedes 5’- and 3’-end processing

In yeast, tRNA splicing occurs in the cytoplasm with tRNAs that underwent 5’- and 3’-end processing (so, the end maturation precedes splicing) (25-27). In vertebrates, using microinjection studies into nucleus of Xenopus oocytes (12,28-30), it was suggested that tRNA splicing is nuclear and splicing precedes the tRNA end processing (reviewed in (11)). It should be noted however that oocytes are non-dividing specialized cells, which are arrested in G2 phase of meiosis I. Because of the specific nature of oocytes, they are loaded with diverse RNA species and do not actively transcribe RNAs until entrance into the other phases of cell cycle and may have also important differences in tRNA transcription/processing when compared to actively divided cells that are dependent on tRNA production. As principal enzymes for both tRNA splicing, ligation and end processing are known in human cells, we decided to use RNAi to determine effects of protein down-regulation on tRNA maturation.

Using siRNAs against RTCB, and its co-factors DDX1 and ARCH (archease) (efficiency of knockdown is shown in Figure 3A), we examined cytoplasmic pre-tRNAs in control versus tRNA ligase complex-depleted cells using NB. Knockdown of RTCB, ARCH and DDX1 promotes formation of 5’-tRNA halves (5’-tRNA exons) that contain 5’-leader sequences in both tRNA^Tyr-GTA^ and tRNA^Leu-CAA^ (Figure 3B, Supplementary Figure S5A). In addition, using NB probes against 3’-tRNA exon, we observe accumulation of 3’-tRNA exon of tRNA^Tyr-GTA^ together with 3’-trailer (Supplementary Figure S5B). We can also conclude that these tRNA fragments do not contain intron sequences. It was recently reported that high-dose H_2_O_2_ treatment of the cells induce oxidative inactivation of RTCB ligase complex (31). As expected, H_2_O_2_ treatment induced accumulation of 5’-leader-exon and 3’-exon-trailer fragments like RTCB knockdown (Supplementary Figure S6). Further, we depleted other components of tRNA splicing machinery, TSEN2 (catalytic subunit of TSEN complex) and CLP1 (an RNA kinase that is hypothesized to participate in a yeast-like splicing/ligation pathway in human cells). Efficient knockdown of TSEN2 and CLP1 (Figure 3C), promotes accumulation of the most immature (end-extended, intron-containing) pre-tRNA^Tyr^ in the cytoplasmic fraction (Figure 3D and Supplementary Figure S5C), which is theoretically conceivable because inhibition of TSEN complex should inhibit the endonucleolytic removal of exons. In addition, both TSEN2 and CLP1 knockdown induced an additional band above that for mature tRNA (Figure 3D: probe 1-34). This molecule contained an intron sequence (Figure 3D: Intron probe), while it did not contain 5’-leader (Figure 3D: 5’-Leader probe), suggesting that this RNA is intron-containing, but end-processed pre-tRNA^Tyr^ (Supplementary Figure S5D). These data suggest that end-processing can take place under the condition where TSEN complex is inhibited. Taken together, based on these data of the effects of the inhibition of tRNA splicing/ligation machinery, these data suggest the model of cytoplasmic pre-tRNA processing where tRNA splicing precedes 5’- and 3’-end maturation in human cells (Figure 3E).

**Figure 3.**
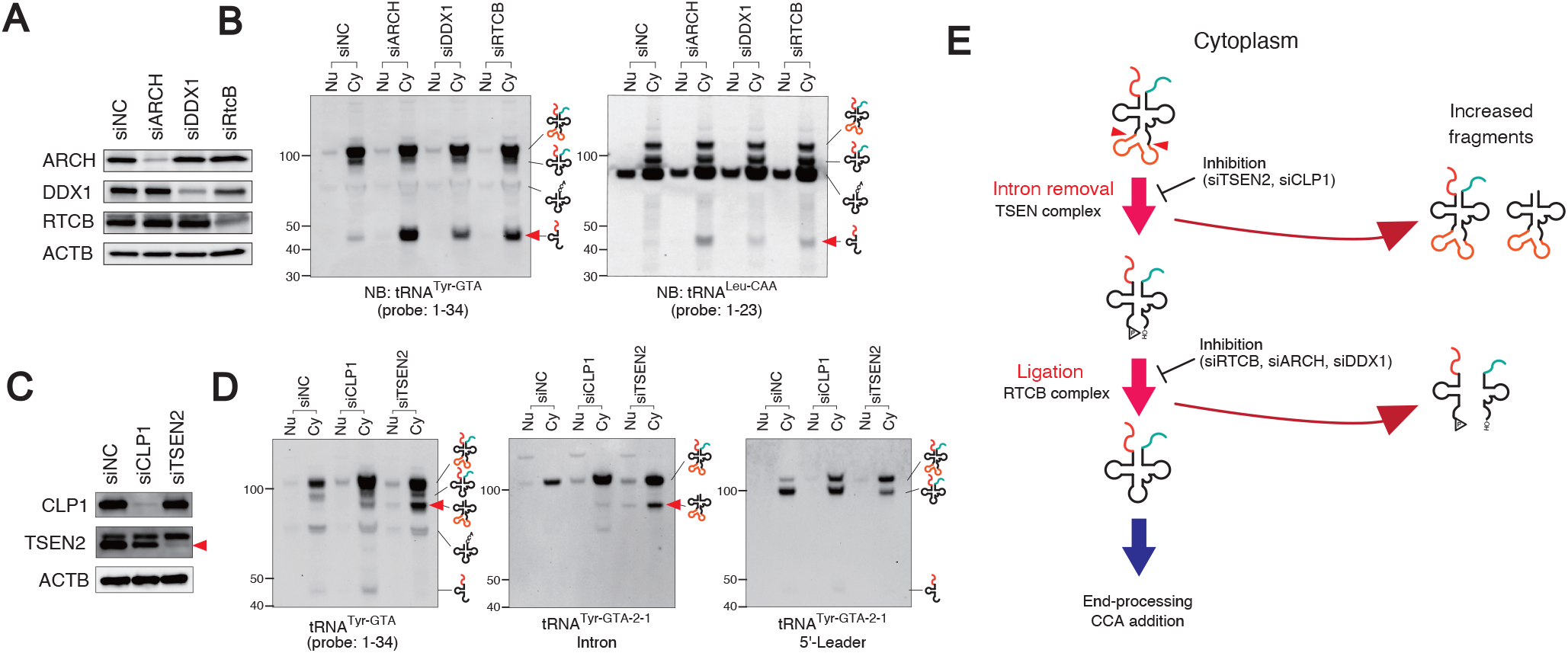
tRNA splicing takes place in the cytoplasm and precedes 5’- and 3’-end processing. (A-B) Knockdown of tRNA ligase components induces 5’-leader-exon fragment in the cytoplasmic fraction. (A) siRNA-mediated knockdown of the components of RTCB ligase complex. (B) Northern blotting for 2 intron-containing tRNAs. Red arrowhead indicates 5’-leader-exon fragments induced by the inhibition of RTCB ligase complex. (C-D) Knockdown-mediated inhibition of TSEN complex induces the accumulation of intron-possessing pre-tRNA^Tyr-GTA^-derived fragments in the cytoplasmic fraction. (C) siRNA-mediated knockdown of CLP1 and TSEN2. Red arrowheads indicate the band for TSEN2. (D) Northern blotting using three different (specific to 5’-exon, intron and 5’-leader) probes targeting tRNA^Tyr-GTA^. Red arrowheads indicate the abnormal (intron-containing, but end-processed) fragment induced by the inhibition of TSEN complex. (E) Schematic diagram of the effect of the inhibition of tRNA splicing component. Inhibition of TSEN complex induces the accumulation of intron-containing (both end-possessing and end-processed) fragments in the cytoplasmic fraction. On the other hand, inhibition of RTCB ligase complex induces end-processed fragments (e.g. 5’-leader-exon and 3’-exon-trailer) in the cytoplasmic fraction, which shows that splicing precedes end-processing in human cells.

### Cytoplasmic La is bound to pre-tRNAs

La protein (autoantigen La, also known as SSB) has been implicated in several processes, the most ubiquitous of which is the binding to and stabilization of newly synthesized Pol III transcripts, including pre-tRNAs (32,33). Binding occurs via the common UUU-OH 3’-terminal motif which results from transcription termination within the Pol III termination signal, oligo(dT) (34). La protein is reported to be both nuclear and cytoplasmic (35), where it binds to subclass of mRNAs bearing pyrimidine-rich TOP motif (36,37). We reasoned that because tRNA processing (namely splicing, ligation, end processing and, possibly CCA-addition) occurs in the cytoplasm, nuclear La may assist to export of pre-tRNA into cytoplasm via binding to UUU-OH 3’-trailer.

To determine whether La binds to pre-tRNA in the cytoplasm, we first fractionated U2OS lysates into nuclear and cytoplasmic fractions (Figure 4A) and then used antibodies against endogenous La protein for immunoprecipitation (IP). As it is seen in Figure 4B, we successfully immunoprecipitated La protein from the cytoplasmic fraction. Our data suggest that La also binds many different RNA species in the cytoplasm (Figure 4C), including intron-containing (tRNA^Tyr-GTA^ and tRNA^Leu-CCA^) and intron-less (tRNA^iMet-CAT^ and tRNA^Gly-GCC^) pre-tRNAs (Figure 4D). The same results were obtained when performing RNA-IP using the cytoplasmic lysate prepared by hypotonic buffer-based fractionation method like in Figure 1-3 (Supplementary Figure S6). Interestingly, we could detect pre-tRNA molecules that contain intron, 5’-leader and 3’-trailer as well as spliced, 5’-leader/3’-end containing pre-tRNA species bound to La protein (Figure 4D), which suggests that La protein keeps binding to pre-tRNAs during splicing. This result is consistent with the finding that splicing precedes end-processing shown in Figure 3. Moreover, components of TSEN complex are also found in the complex with La (Figure 4E) in contrast to the proteins suggested to be involved in primary tRNA export (XPOT, XPO5 and XPO1). La binding to TSEN complex is RNase-sensitive (Figure 4F), and specifically dependent on the levels of pre-tRNAs (Figure 4G, H). Treatment of cells with RNase A or transcription inhibitor Actinomycin D (ActD) also enhances IF signals of TSEN2 in the cytoplasm (Figure 4I, J). These data suggest that La may be interacted with TSEN2 complex through pre-tRNAs in the cytoplasm. This interaction might inhibit the binding of TSEN2 antibody to TSEN2 when performing IF (Figure 4K).

**Figure 4.**
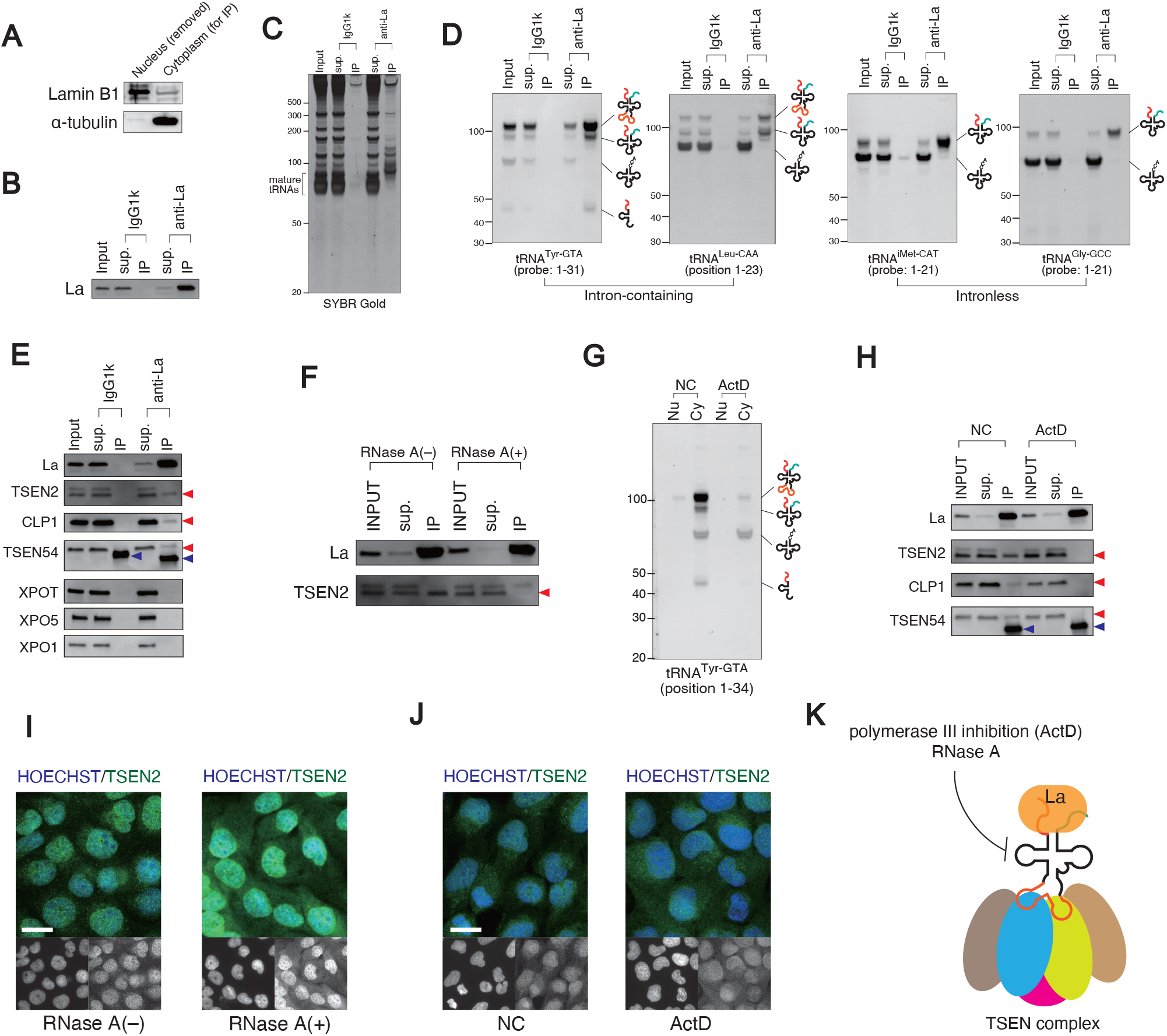
Cytoplasmic La is bound to pre-tRNAs and keeps binding during splicing. (A-D) Cytoplasmic fraction contains La-bound pre-tRNAs. (A) Cytoplasmic fraction was subjected to following RNA-IP analysis using anti-La antibody. (B) Validation of immunoprecipitation. La protein was efficiently pulled down by anti-La antibody. (C) SYBR Gold staining of purified RNAs derived from IP fraction immunoprecipitated by anti-La antibody. (D) Northern blotting for both intron-containing (Tyr-GTA and Leu-CAA) and intron-less (iMet-CAT and Gly-GCC) tRNAs. Note that pre-tRNAs both before and after splicing were pulled down in intron-containing genes, suggesting that La keeps binding to pre-tRNAs throughout splicing. (E-J) La-bound pre-tRNAs are interacted with TSEN complex in the cytoplasm. (E) TSEN components are co-immunoprecipitated with La, while exportins not. TSEN components co-immunoprecipitated with La were indicated by red arrowheads. Bands detected below TSEN54 band (indicated by blue arrowheads) are antibodies used by IP (mouse IgG1k or anti-La antibody). (f) Pre-treatment of the lysate with RNase A decreases binding of TSEN2 to La, suggesting that TSEN complex is interacted with La through pre-tRNAs. (G) Actinomycin D (ActD) treatment (at 5 µg/ml for 2 hr) decreased the amount of pre-tRNAs, while mature tRNA was not changed. (H) Under the condition, ActD treatment abolished the interaction between La and TSEN components (indicated by red arrowheads). Bands detected below TSEN54 band (indicated by blue arrowheads) are antibodies used by IP (mouse IgG1k or anti-La antibody). (I) Cytoplasmic TSEN2 signal is increased by RNase A treatment. After fixation and permeabilization, samples were treated with 100 µg/ml bovine RNase A at 37 °C for 1 hr. (J) Cytoplasmic TSEN2 signal is also increased by ActD treatment. Cells were treated with 5 µg/ml ActD for 3 hr before fixation. (K) Schematic model of the interaction between La and TSEN complex in the cytoplasm. La is interacted with TSEN complex in the cytoplasm through pre-tRNAs. This interaction can inhibit the binding of anti-TSEN antibodies to TSEN complex when performing IF, which is possible to cause decrease in the cytoplasmic signals.

La appears to be freely soluble and readily distributes between nuclear and cytoplasmic fractions during cell disruption for microscopy and biochemical preparations (discussed in (38)). Alternative methods were used to demonstrate its cytoplasmic localization and association with pre-tRNAs (Supplementary Figure S8-S10). Moreover, we tested whether in response to stress, La-bound pre-tRNAs are located into Stress Granules (SGs), cytoplasmic RNA granules with pro-survival properties that regulate several aspects of RNA metabolism (39). Both La protein (IF with three independent La-specific antibodies) and pre-tRNAs (three different species) are associated with SGs (Supplementary Figure S11) suggesting that La-bound pre-tRNAs are found in the cytoplasm. In summary, we conclude that cytoplasmic La keeps binding to pre-tRNAs throughout splicing and ligation (Figure 4K).

### Cytoplasmic La is involved in nuclear-cytoplasmic export of pre-tRNAs

Although tRNAs are small molecules, exit of tRNAs from the nucleus is not a passive process. Members of the conserved β-importin superfamily are primarily candidates for the role on transporting pre-tRNA to the cytoplasm. Members of this superfamily utilize the RAN (Ras-related nuclear protein) pathway, which relies on the action of the small GTPase RAN. RAN is primarily found in GDP bound form in the cytoplasm and in the GTP bound form in the nucleus (40).

In vertebrates, XPO5 (Exportin-5), XPOT (exportin-t) and XPO1 (Crm1) are acting via RAN-dependent pathway and proposed to be main players in the primary tRNA nuclear export from the nucleus to the cytoplasm (reviewed in (10)). However, it should be noted that genes are scored as unessential in different genome-wide screen studies (41-43) proposing an existence of alternative or redundant pathways for the primary nuclear export. In attempt to characterize machinery that export pre-tRNAs to the cytoplasm we depleted either RAN itself, or RAN-dependent tRNA exporters XPOT, XPO5, XPO1 in U2OS cells by shRNAs (Supplementary Figure S12, S13). We then obtained nuclear and cytoplasmic fractions from control and depleted cells and determined effects of their knockdown on the distribution of pre-tRNAs (Supplementary Figure S12, S13). Our data suggest that efficient depletion of RAN and XPO1/XPO5/XPOT does not affect export of pre-tRNA into the cytoplasm. In addition, treatment with leptomycin B, the specific inhibitor of XPO1 (44), also did not affect localization of pre-tRNAs (Supplementary Figure S13D-F). Finally, depletion of NXF1, the factor that was shown to bind tRNAs and affect tRNA export in RAN-independent manner in yeast (45), also did not affect pre-tRNA distribution (Supplementary Figure S14). These data suggest that pre-tRNA export in human cells is different from tRNA export in lower eukaryotes.

Our data suggest that cytoplasmic La is bound to pre-tRNAs. As La protein found both in the nucleus and the cytoplasm, we hypothesized that La may directly assist on some aspects of pre-tRNA export from the nucleus to the cytoplasm. To determine such role(s), we employed shRNA-mediated depletion of La in U2OS cells (efficiency of knockdown is shown in Figure 5A). We then fractionated control and La-depleted cell lysates into the nuclear and cytoplasmic fractions, and isolated total RNA from both fractions. Levels of nuclear and cytoplasmic pre-tRNA species (both intron-containing tRNA^Tyr-GTA^ and intron-less tRNA^iMet-CAT^) were then compared using NB analysis and quantified (three biological replicates, Figure 5B, C). Knockdown of La protein does not affect levels of mature tRNAs (Figure 5B, Supplementary Figure S15). However, La knockdown induced the accumulation of end-extended pre-tRNAs both with and without intron in the nuclear fraction (Figure 5B, C and Supplementary Figure S15), which suggests that La is actively involved in pre-tRNA export from the nucleus to the cytoplasm.

**Figure 5.**
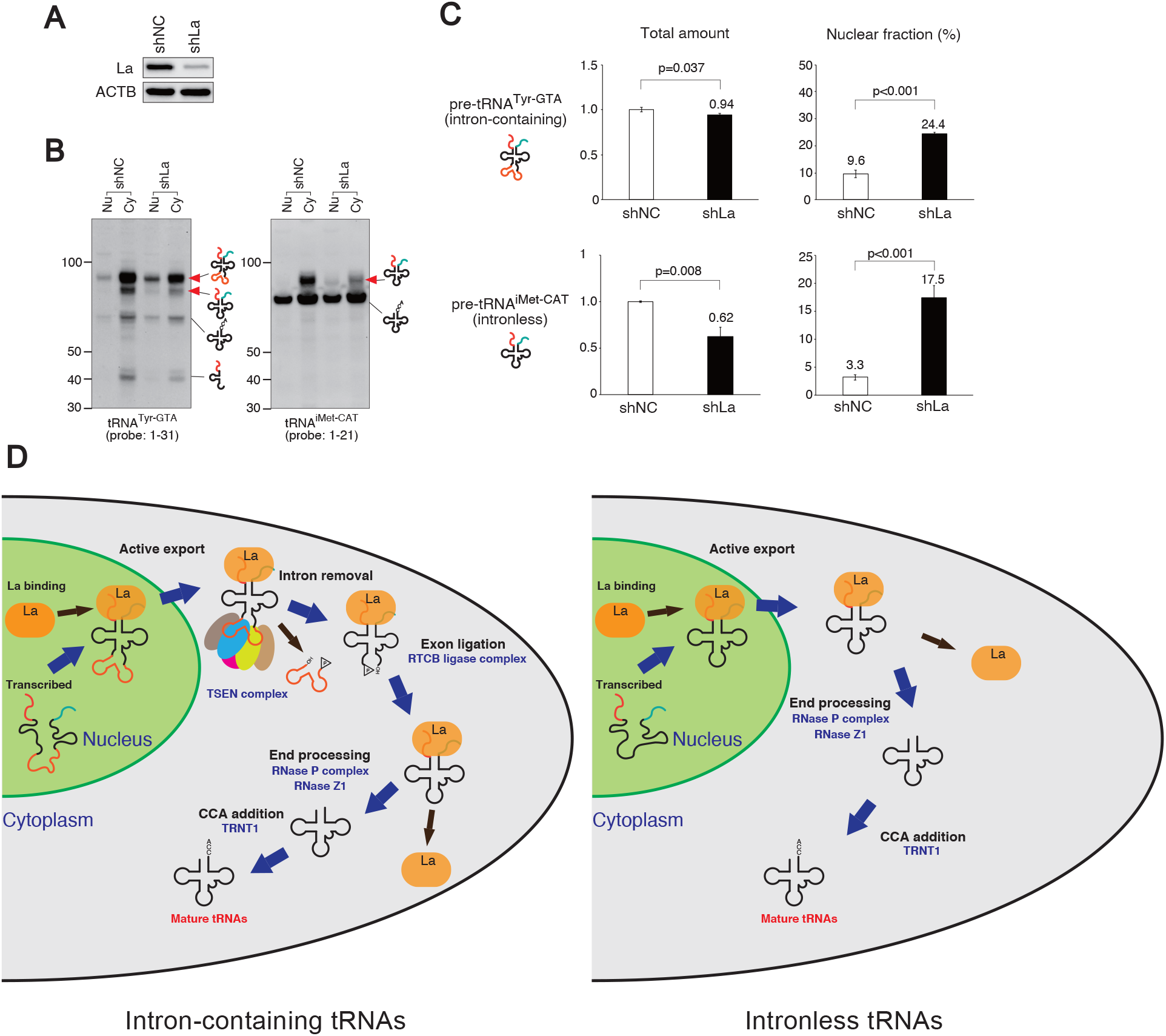
La protein is involved with pre-tRNA export to the cytoplasm. (A) shRNA-mediated knockdown of La. (B-C) La knockdown induces accumulation of end-extended pre-tRNAs in the nuclear fraction. (B) Northern blotting for intron-containing (Tyr-GTA) and intron-less (iMet-CAT) tRNAs. Red arrowheads indicate pre-tRNAs affected by La knockdown. (C) Densitometry analysis of the pre-tRNA signals shown in (B). Data are presented as the mean ± SD of triplicate independent experiments. P-values for Student’s t-test are shown. (D) Schematic model of alternative tRNA maturation pathway in human cells. First, transcribed pre-tRNAs are exported into the cytoplasm as a complex with La protein. Next, pre-tRNAs are spliced in the cytoplasm before end-processing. During splicing, La remains binding to pre-tRNAs. Then end-processing and CCA-addition take place in the cytoplasm.

## DISCUSSION

Pioneer work on the determination of the intracellular localization of pre-tRNAs in mammalian cells suggested that these precursor molecules are found in the cytoplasmic fractions (46-49). It was suggested that “during the maturation process leading to the formation of biologically functional transfer RNA, the precursor tRNA molecules in the cytoplasm might adopt a certain conformation whilst for example methylation of specific bases, or other modifications to the primary structure, are carried out by specific enzymes, such as tRNA methylases, located in the cytoplasm” (48,49). Further, conversion of pre-tRNA into tRNA has been shown *in vitro* using crude HeLa cytoplasmic extracts (50,51) and HeLa S100 cell-free extracts (52). Subsequently, it was shown that cloned tRNA genes microinjected into the nucleus of frog oocytes can be efficiently transcribed and processed (30). Use of microinjection into “living test tubes” has been very instrumental and served as a decisive step to postulate that tRNA processing in eukaryotes is entirely nuclear event.

Later, data in yeast convincingly showed that some steps of pre-tRNA processing such as pre-tRNA splicing occur in the cytoplasm where the factors involved in tRNA splicing reside (25,27,53). Reminiscent of *S. cerevisiae* and *S. pombe*, protist *Trypanasoma brucei* also localizes components of tRNA splicing machinery in the cytoplasm (54).

Here, by combination of cell fractionation analysis and northern blotting, we demonstrated that unprocessed forms of pre-tRNAs (both intron-containing and non-intron families) are readily detected in the cytoplasmic fractions of various human cell lines (Figure 1 and Supplementary Figure S1). We further developed pre-tRNA-specific FISH protocol relying on the detection of different parts of unprocessed tRNAs (Figure 1D, and validated in Supplementary Figure S2) that showed cytoplasmic distribution of pre-tRNAs, in agreement with cell fractionation results. Our data are consistent with the recent observation that intron-containing pre-tRNA^Ile^ is enriched in the cytoplasm in human HEK293T cells (55).

The cytoplasmic localization of pre-tRNAs suggested that at least some of their processing steps are cytoplasm-associated events as described in lower eukaryotes. Initially, we focused on the localization of the TSEN complex, which has been reported to be associated with mitochondrial surfaces in yeast. Using cell fractionation approach and western blotting against endogenous proteins, we demonstrated that all four subunits of TSEN complex are readily detected predominantly in the cytoplasmic fraction (Figure 1A). Since this is in contrast with previously reported nuclear localization of TSEN2 and TSEN34 in the nucleus (2), we further validated cytoplasmic localization of catalytic subunit of TSEN complex, TSEN2, by using IF microscopy and shRNA-mediated depletion experiments aimed on the specificity of used antibodies (note that 3 independent shRNA constructs as well as siRNA, and four independent, commercially-available antibodies were used (Supplementary Figures S3 and S4). Our data suggest that endogenous TSEN2 localizes in the cytoplasm in contrast to exclusively nucleus-localized over-expressed recombinant GFP-tagged proteins that were used in Paushkin et al. studies. It should also be noted, that in yeast, over-expression of tagged Sen subunits can cause subcellular mislocalization (27), thus reinforcing the importance of the detection of proteins in endogenous settings. To the best of our knowledge, our results are the first evidence that endogenous TSEN subunits are localized dominantly in the cytoplasm.

Besides TSEN splicing endonuclease complex, CLP1 kinase is another factor that is implicated in pre-tRNA splicing (56). Our nuclear-cytoplasmic fractionation and IF experiments showed that CLP1 localizes to both compartments (Figure 2), which is also in agreement with nuclear role of CLP1 in pre-mRNA processing (57). Our data also shows that CLP1 knockdown inhibits intron cleavage similarly to TSEN2 knockdown (Figure 3D) suggesting that CLP1 is also essential factor for intron cleavage *in cellulo*, although previously published in vitro-based data suggested that CLP1 is not required for intron cleavage (20).

Further analysis also demonstrated (Figure 2) that: 1) RtcB and DDX1 components of tRNA splicing ligase complex are localized in both nucleus and cytoplasm while the essential subunit Archease is cytoplasmic; 2) components of RNase P complex required for 5’-leader processing (POP4/Rpp21 and H1 RNA) are largely cytoplasmic; 3) RNase Z1, required for 3’-trailer removal, is cytoplasmic; 4) CCA-adding enzyme TRNT1 localizes to the cytosol. RTCB ligase complex is suggested to function in the cytoplasm because one of the essential subunits Archease is exclusively localized in the cytoplasm (58). Our fractionation data are consistent with this report. In addition, another essential subunit PYROXD1, which is cytosolic, was recently identified (31), further suggesting that RTCB ligase complex functions predominantly in the cytoplasm. For RNase P complex, although its protein subunits POP4 and Rpp21 are present in both nuclear and cytoplasmic fraction, its RNA component H1 RNA is highly enriched in the cytoplasmic fraction, suggesting that RNase P complex may function predominantly in the cytoplasm. POP4 is suggested to be shared with RNase MRP complex which is believed to function in the nucleus, while Rpp21 is thought to be RNase P-specific (59), which might be able to explain why POP4 localizes in both fraction and Rpp21 is cytoplasm-dominant. Taken together with results of pre-tRNA and TSEN complex localization studies, these data suggest that both pre-tRNAs and tRNA processing machineries co-exist in the cytoplasm.

siRNA depletion against components of tRNA splicing machineries TSEN2 and CLP1 and RTCB ligase complex components RTCB, DDX1 and ARCH allowed us to determine the order of pre-tRNA processing in the cytoplasm (Figure 3 and Supplementary Figure S5 and S6). Based on the detection of the pre-tRNA intermediates using northern blotting analysis of intron-containing tRNAs^Tyr/Leu^, we conclude that intron removal precedes 5’- and 3’-end processing in human cells (Figure 3E). From one side, our data agrees with previously published data that in vertebrates 5’-3’ maturation occurs after tRNA removal (29), but in the cytoplasm and not in the nucleus. From the other side, we agree with data from lower eukaryotes that tRNA processing takes place in the cytoplasm, however in yeast 5’- and 3’-end processing happens before tRNA intron removal (reviewed in (15)).

Our data also suggest that upon transcription by RNA Pol III, tRNA precursors associate with La protein and export from the nucleus to the cytoplasm as pre-tRNA/La complexes. Cytoplasmic La binds both intron-less and intron-containing pre-tRNAs. Interestingly, La remains bound to end-extended pre-tRNAs both with and without introns (Figure 4D), indicating that La protein continues to be associated with pre-tRNAs during splicing. This result is also in agreement with results showing that splicing precedes end-processing (Figure 3E).

Another important aspect of our findings is discrepancy between IF and subcellular fractionation data, especially the localization of La. Human La protein is known to be mostly (∼80%) nuclear (reviewed in (32)). Our IF result shows that La localizes exclusively in the nucleus (Supplementary Figure S9), while subcellular fractionation shows that La is predominantly present in the cytoplasmic fraction (Supplementary Figure S8D and S13D). Supplementary Figure S9 shows that permeabilization step causes massive loss of soluble molecules including tRNAs and pre-tRNA-bound La. Only minor proportions of them remained in the fixed sample. This data suggests that IF/FISH does not reflect the actual amount of soluble molecules, or nucleus/cytoplasm ratio of them. We believe that one of the reasons for the discrepancy between IF/FISH and subcellular fractionation can be explained by the loss of soluble molecules during sample preparation for IF/FISH. La is known to be freely soluble and upon cell disruption for microscopy and biochemical preparations (discussed in (38)), it readily distributes between nuclear and cytoplasmic fractions. We used other approaches to demonstrate its cytoplasmic localization and association with pre-tRNAs (Supplementary Figure S8-S10). In addition, La-bound pre-tRNAs are located into cytoplasmic SGs upon sodium arsenite stress (Supplementary Figure S11), thus further proving that La-bound pre-tRNAs are found in the cytoplasm.

Finally, we found that pre-tRNA export in human cells is significantly different from tRNA export in yeast. An efficient depletion of RAN, XPO1/XPO5/XPOT and NXF1 does not impact export of pre-tRNA into the cytoplasm as well as treatment with leptomycin B, the specific inhibitor of XPO1 (Supplementary Figure S13D-F and S14). Instead, our data supports an active or at least supportive role for La protein in pre-tRNA export from the nucleus. Upon depletion of La (Figure 5A), we observe retention of end-extended pre-tRNAs both with and without introns in the nuclear fraction (Figure 5 C) without any effect on the levels of mature tRNAs (Figure 5B, Supplementary Figure S15). Our data suggest that La binds pre-tRNAs in the nucleus and further contributes into pre-tRNA export from the nucleus to the cytoplasm. In addition, the discrepancy of the localization of pre-tRNA processing between microinjection studies (exclusively in the nucleus) and our current study might also be explained by the binding between pre-tRNAs and La in endogenous tRNA maturation pathway.

In summary, our data suggest a model where processing of tRNA precursors is cytoplasmic similarly to the observed in lower eukaryotes, although tRNA splicing precedes 5’- and 3’-end processing in human cells unlike in yeast. Our data also suggest that many aspects of pre-tRNA export in vertebrates maybe significantly different from lower eukaryotes, which warrants future investigation in this area of tRNA biology.

## SUPPLEMENTARY DATA

Supplementary Data are available online.

## ACKNOWLEDGEMENT

We thank members of Ivanov and Anderson labs for discussion and helpful critique.

## FUNDING

This work was supported by the National Institutes of Health [R35 GM126901 to P.A., RO1 GM126150 to P.I., K99 GM124458 to S.M.L.], the Japan Society for the Promotion of Science, Grants-in-Aid for Scientific Research (PS KAKENHI) [26860094 and 21K06865 to Y.A., 19H03509 to Y.H., and 18H02822, 20K20604 and 21H02932 to T.A.], and the Japan Agency for Medical Research and Development (AMED) [20ek0210133h0001, 20ak0101127h0001 and J21000294 to T.A.].

## CONFLICT OF INTEREST

The authors declare no conflict of interest.

## Notes

### Competing Interest Statement

The authors have declared no competing interest.

