## Supplementary Info for "Cytoplasmic processing of human transfer RNAs"

#### Supplementary Method

##### Subcellular fractionation using digitonin buffer or detergent-free hypotonic lysis buffer (related to Supplementary Figure S8)

The cytosolic fraction was obtained by incubating cells with Digitonin buffer [20 mM HEPES (pH 7.0), 150 mM NaCl, 25 µg/ml digitonin (Sigma-Aldrich)] or HLB without NP-40 (detergent-free HLB) on ice for 10 min. After collecting the supernatant as a "Cytosol" fraction, cell pellet was incubated with the regular HLB (containing NP-40) at room temperature for 5 min, then the supernatant was collected as "Organelle" fraction. After washing twice with detergent-free HLB, the nuclear pellet was collected as "Nucleus" fraction.

##### Determination of the amount of leaked cellular content during IF/FISH (related to Supplementary Figure S9)

U2OS cells were seeded onto 6 well plate, then incubated overnight. After fixation with 4% paraformaldehyde for 15 min in the same way as that for IF/FISH, "Total" fraction was obtained by directly adding Trizol (for RNA) or RIPA buffer (for protein) without permeabilization. After permeabilization, the permeabilization buffer was collected as "Leaked" fraction, while the remaining cells was collected as "Fixed" fraction. Cross-linking was reversed by incubating at 65ºC for 1 hr. Note that "Fixed" fraction reflects the molecules evaluated by IF/FISH, while molecules in "Leaked" fraction is never detected by IF/FISH because it is removed from cells during sample preparation. RNA and protein samples from each fraction were subjected to Northern blotting and Western blotting, respectively. In addition, RNA-IP was also performed using the "Leaked" fraction to examine how much La-bound pre-tRNAs was lost from IF/FISH samples.

##### PFA fixation-combined fractionation (related to Supplementary Figure S10)

The cytosolic fraction was obtained by digitonin-based subcellular fractionation after PFA fixation to inhibit active nuclear-cytoplasmic transport for minimizing the leakage of nuclear content during fractionation. After washing cell pellet twice with digitonin-containing buffer, the cell pellet was collected as "The others" fraction. Cross-linking was reversed by incubating at 65ºC for 1 hr. Note that "The others" fraction contains not only nuclear content but also non-soluble content in the cytoplasm including organelles.

Supplementary Figure S1

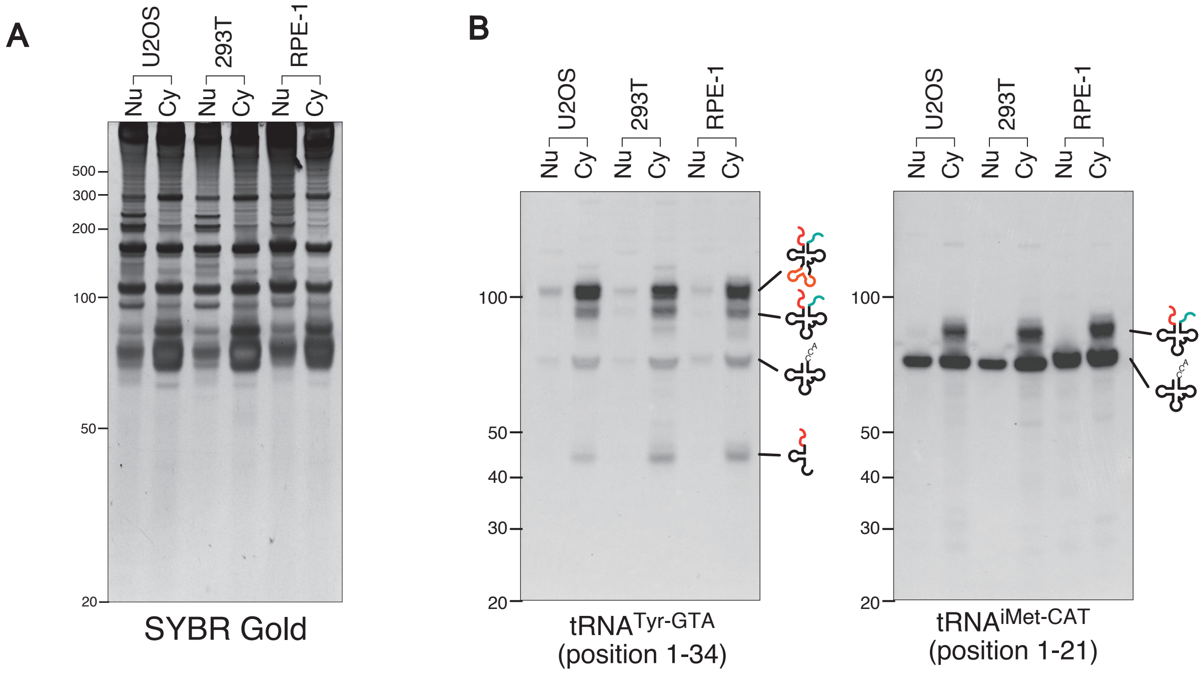


**Supplementary Figure S1.** Other cell lines than U2OS also shows enrichment of pre-tRNAs in the cytoplasmic fraction. (A) SYBR Gold staining of fractionated RNAs derived from U2OS, 293T and RPE-1 cells. (B) Northern blotting shows enrichment of pre-tRNAs in the cytoplasmic fraction regardless of cell line.

### Supplementary Figure S2

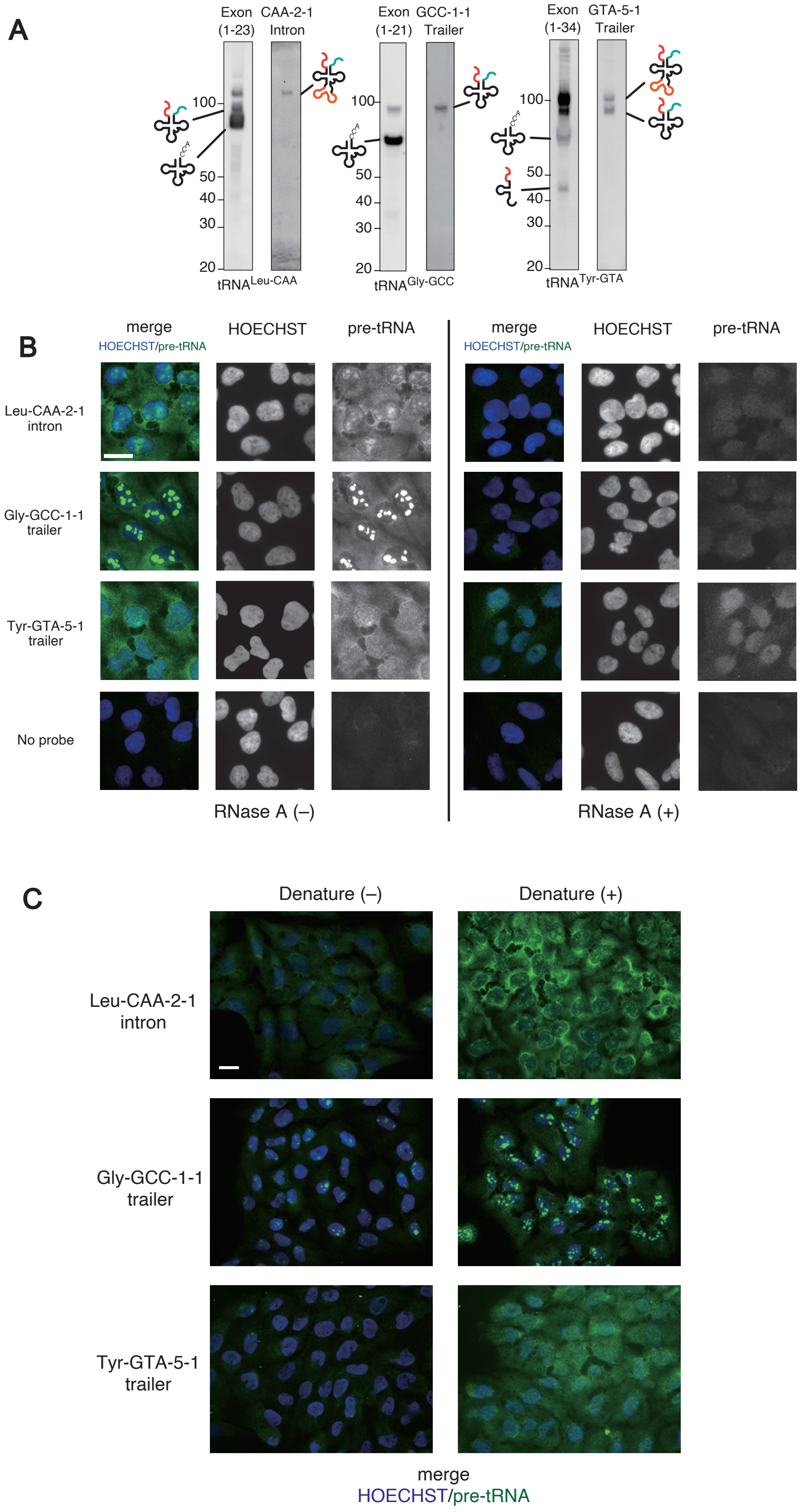

Supplementary Figure S2 (continued)

D

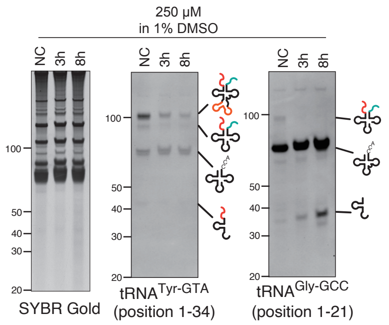


E

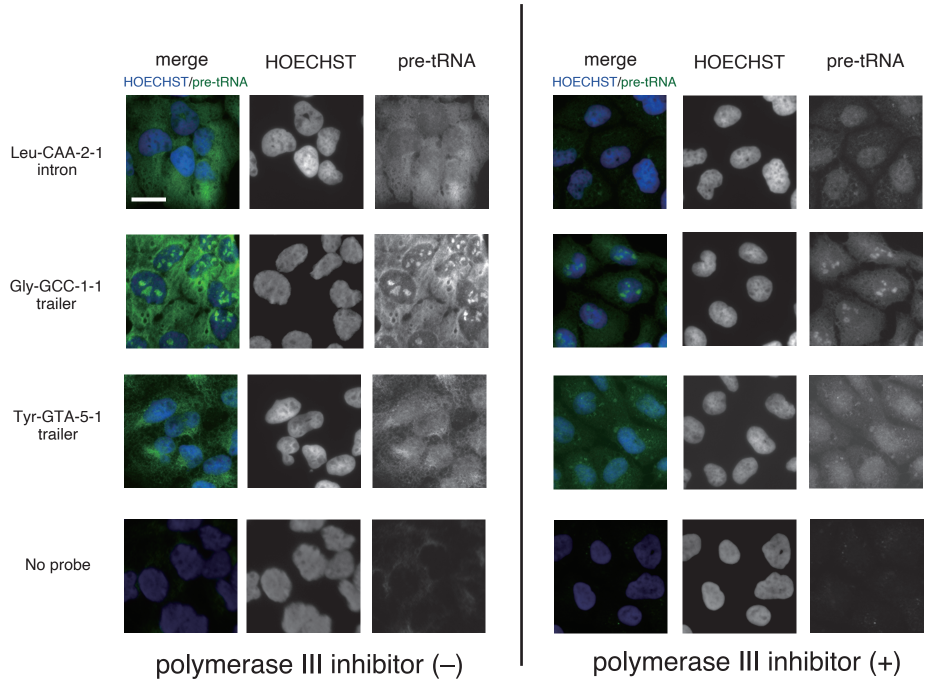


**Supplementary Figure S2.** Validation of pre-tRNA FISH. (A) The probes used for FISH detect only pre-tRNAs by Northern blotting. (B) FISH probes detect RNA molecules. RNase A treatment decreased the cytoplasmic signal, suggesting that the probes were hybridized to RNA molecules. (C) FISH probes detect folded RNAs. Denature step before hybridization significantly increased the signals, suggesting that probes target folded RNAs. (D-E) Polymerase III inhibitor decreases pre-tRNA signal in both Northern blotting and pre-tRNA FISH.
(D) Northern blotting. Polymerase III inhibitor treatment decreased the amount of pre-tRNAs, while the amount of mature tRNAs did not change. (E) pre-tRNA FISH. Polymerase III inhibitor (557403, Sigma-Aldrich) treatment at 250 µM for 3 hours also decreased the signals.

Supplementary Figure S3

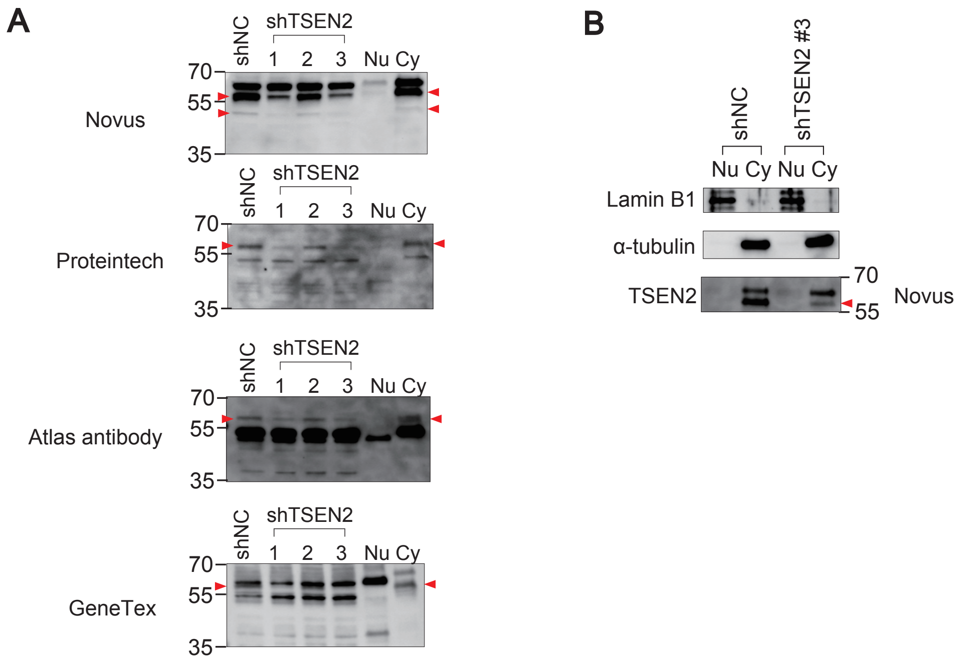


**Supplementary Figure S3.** TSEN2 localizes in the cytoplasm. (A) Three shRNA constructs were generated for TSEN2 knockdown. Construct #1 and #3 worked among them. TSEN2 was detected as a band just above 55 kDa marker (indicated as red arrowheads) in all the 4 antibodies, and it was enriched in the cytoplasmic fraction. Note that Novus antibody also detected an additional band below 55 kDa marker, suggesting an isoform. (B) TSEN2 knockdown does not alter the cytoplasmic localization.

Supplementary Figure S4

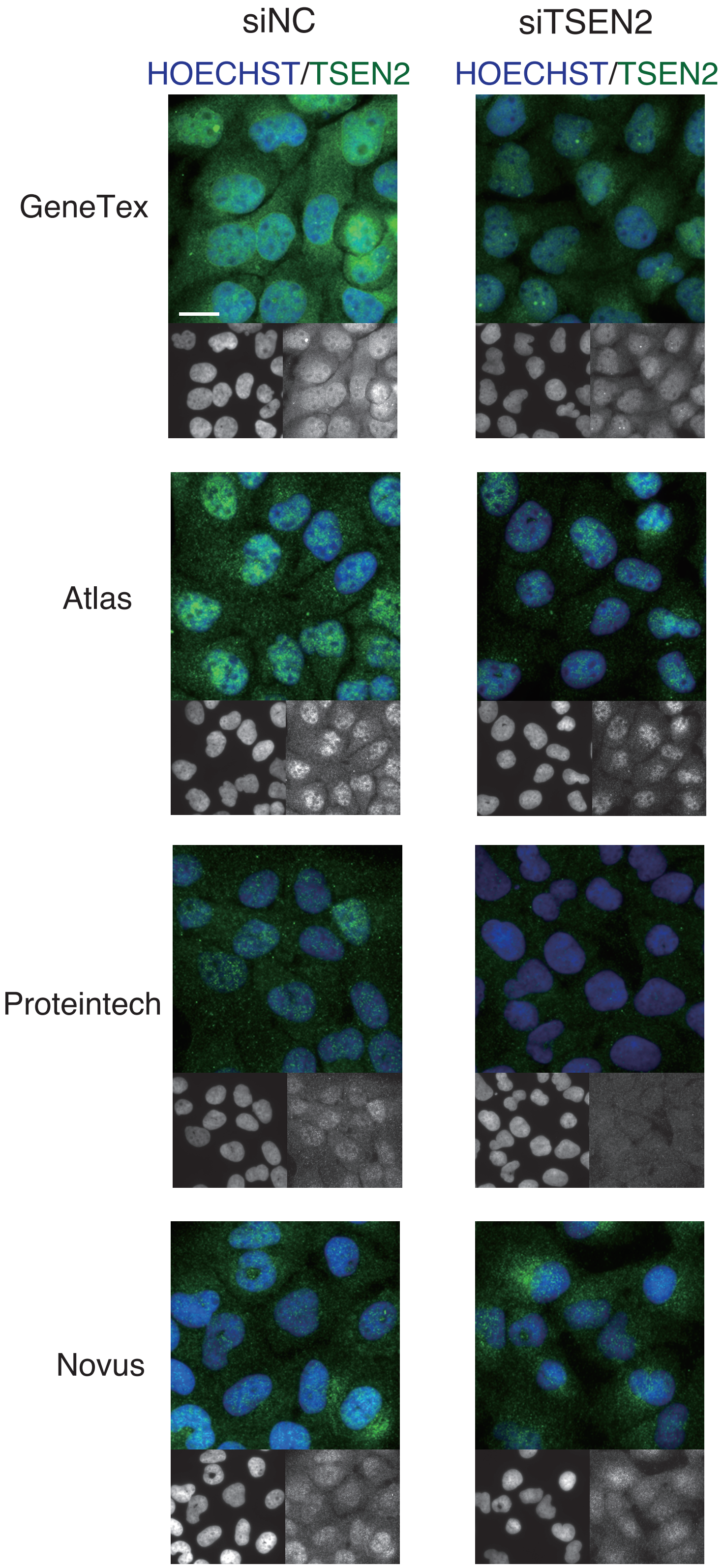


**Supplementary Figure S4.** Immunofluorescence for TSEN2 combined with siRNA-mediated knockdown. The results using four TSEN2-specific antibodies are shown.

### Supplementary Figure S5

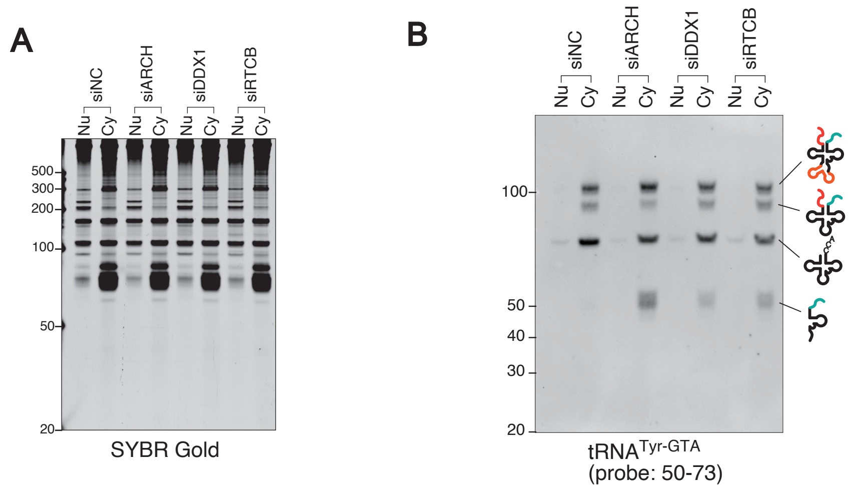


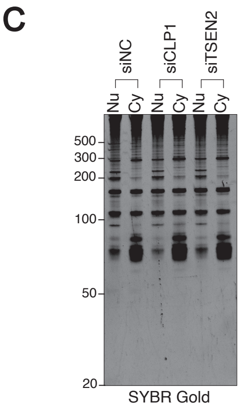


**D**

| tRNA$^{\text{Tyr-GTA-2-1}}$ | Estimated length |
|---|---|
| pre-tRNA with 5'-leader, intron and 3'-trailer | 111 nt |
| pre-tRNA after splicing (with 5'-leader and 3'-trailer) | 95 nt |
| end-processed pre-tRNA (with intron) | 89 nt |
| mature tRNA (with 3'-CCA termini) | 76 nt |

**Supplementary Figure S5.** Additional data for Figure 3. (A) SYBR Gold staining of fractionated RNAs under knockdown of RTCB ligase components. (B) Knockdown of RTCB ligase components induced 3'-exon-trailer fragment as well as 5'-leader exon fragment. (C) SYBR Gold staining of fractionated RNAs under knockdown of TSEN components. (D) Estimated length of tRNA$^{\text{Tyr-GTA-2-1}}$-related molecules.

Supplementary Figure S6

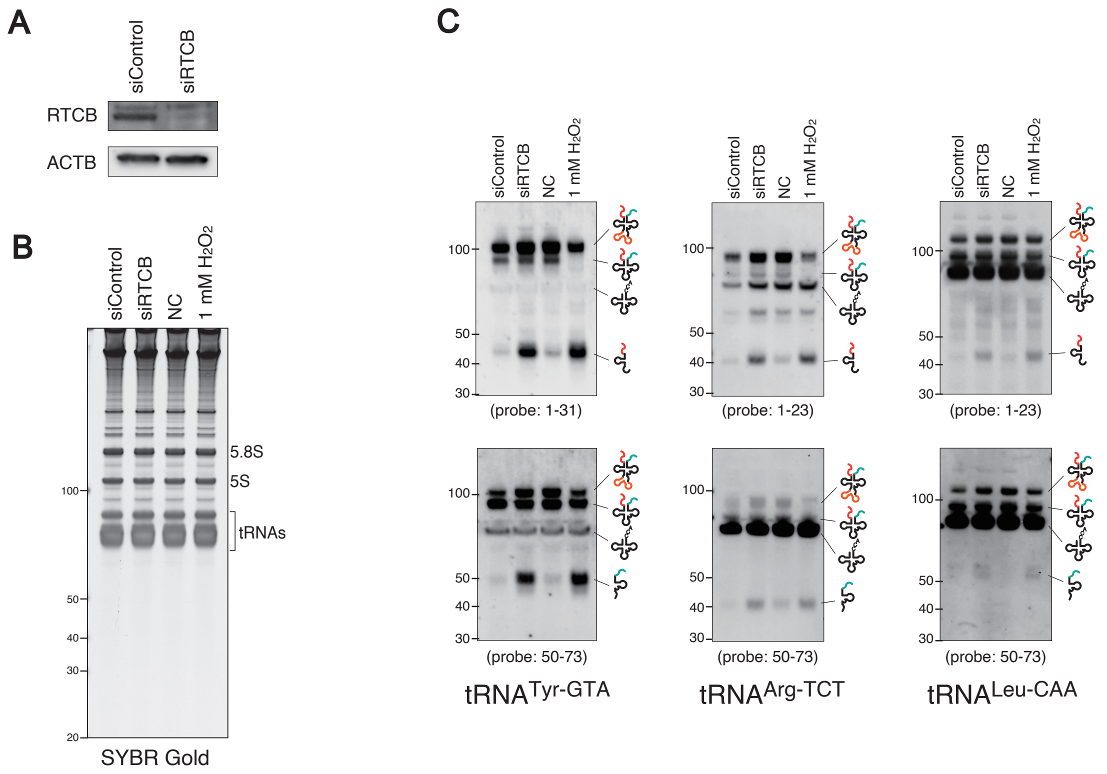


**Supplementary Figure S6.** Both RTCB knockdown and $H_2O_2$ treatment induces 5'-leader-exon and 3'-exon-trailer fragments. (A) siRNA-mediated RTCB knockdown. (B) SYBR Gold staining of total RNAs under RTCB knockdown or $H_2O_2$ treatment. (C) Northern blotting. Both RTCB knockdown and $H_2O_2$ treatment induced 5'-leader-exon and 3'-exon-trailer fragments derived from intron-containing tRNA genes, which suggests that $H_2O_2$ treatment inhibits the activity of RTCB ligase complex as previously indicated.

### Supplementary Figure S7

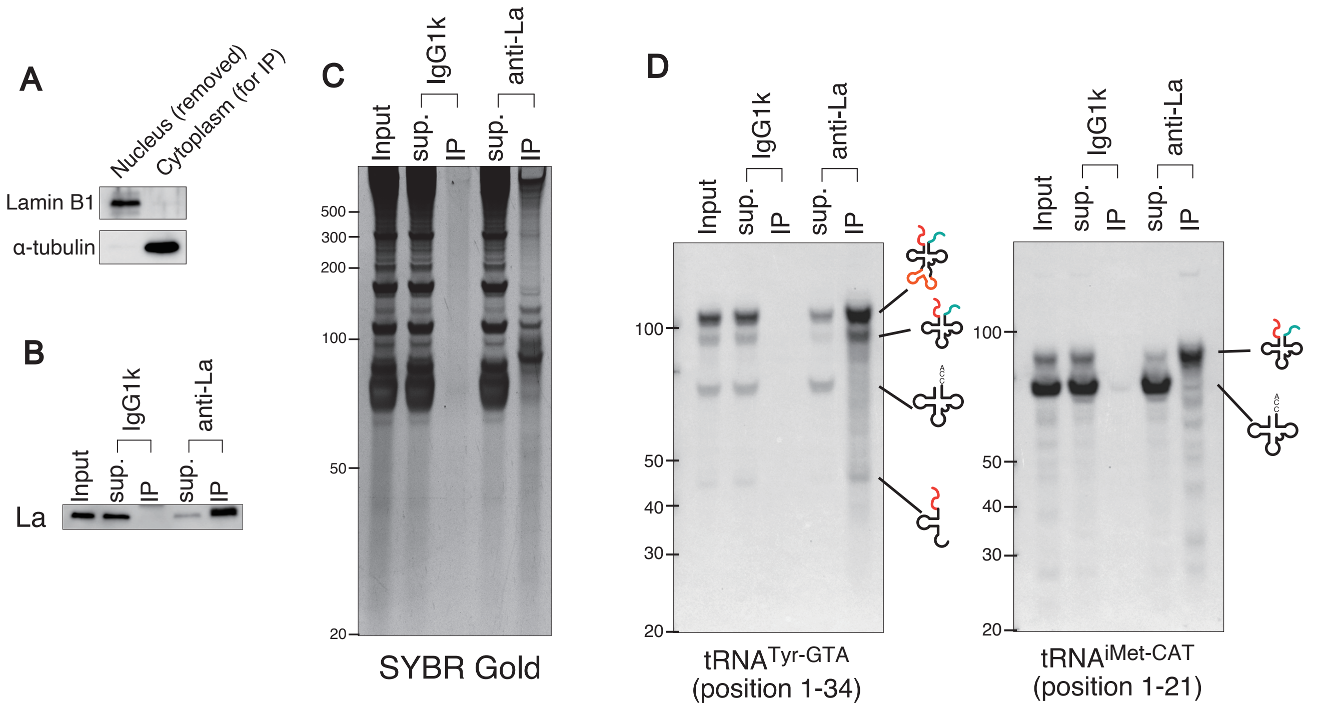


**Supplementary Figure S7.** RNA-IP analysis using the cytoplasmic lysate obtained by hypotonic buffer-based fractionation method in Figure 1-3. (A) Cytoplasmic fraction obtained by hypotonic buffer-based fractionation was subjected to following RNA-IP analysis using anti-La antibody. (B) Validation of immunoprecipitation. La protein was efficiently pulled down by anti-La antibody. (C) SYBR Gold staining of purified RNAs derived from IP fraction immunoprecipitated by anti-La antibody. (D) Northern blotting for both $tRNA^{Tyr\text{-}GTA}$ (intron-containing) and $tRNA^{iMet\text{-}CAT}$ (intron-less).

#### Supplementary Figure S8

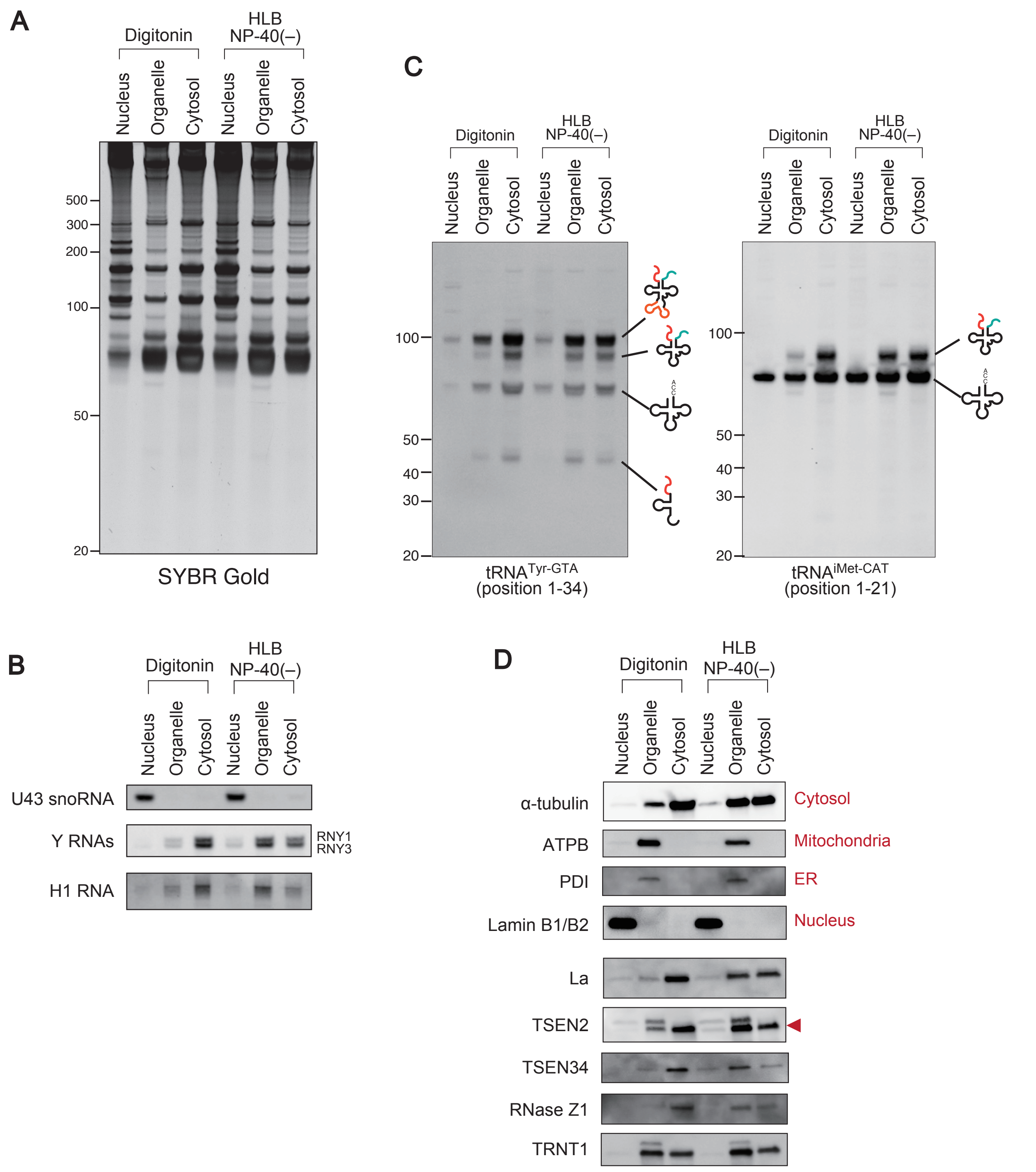


**Supplementary Figure S8.** Cytosolic fraction contains pre-tRNAs, La and other tRNA-processing molecules. Cytosolic fraction was obtained by digitonin-containing buffer or hypotonic lysis buffer without any detergent. (A) SYBR Gold staining of each fraction. (B) Northern blotting for Nuclear RNA (U43 snoRNA) and cytoplasmic (Y-RNAs). Northern blotting for H1 RNA is also shown. (C) Northern blotting for both tRNA$^{Tyr\text{-}GTA}$ (intron-containing) and tRNA$^{iMet\text{-}CAT}$ (intron-less). (D) Western blotting for La and other tRNA-processing molecules.

### Supplementary Figure S9

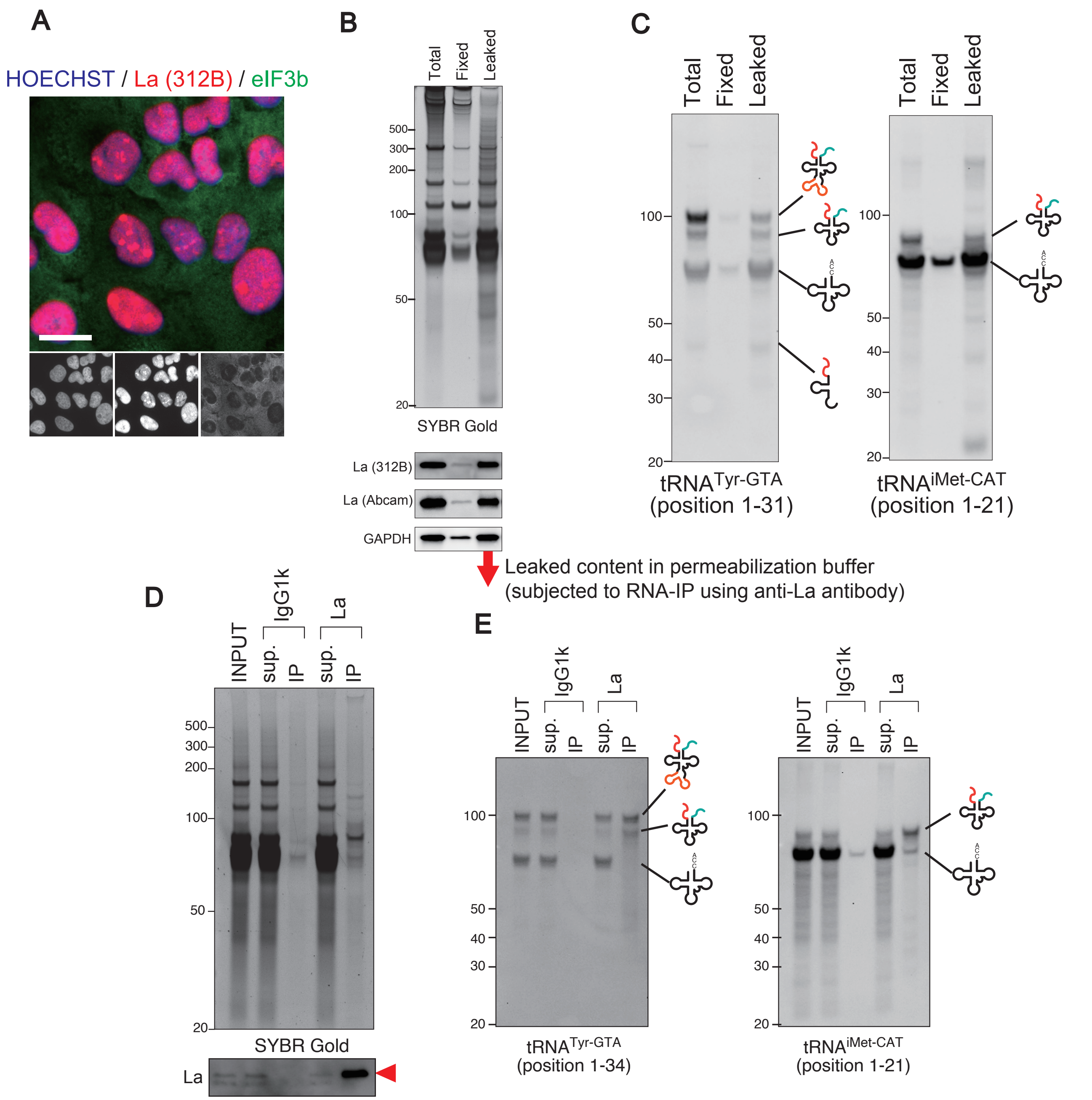


**Supplementary Figure S9.** Immunofluorescence causes massive loss of RNAs including La-bound pre-tRNAs. After fixation with 4% paraformaldehyde and permeabilization, the permeabilization buffer was collected as "Leaked" fraction, while the remaining cells was obtained as "Fixed" fraction. (A) Representative immunofluorescence image of La. (B) La and RNAs are massively lost into permeabilization buffer. SYBR Gold staining for purified RNAs and Western blotting for La are shown. (C) pre-tRNAs are also massively lost during permeabilization. Northern blotting for $tRNA^{Tyr\text{-}GTA}$ (intron-containing) and $tRNA^{iMet\text{-}CAT}$ (intron-less) are shown. (D-E) Leaked fraction contains La-bound pre-tRNAs. RNA-IP using anti-La antibody was performed using leaked fraction collected after permeabilization. (D) SYBR Gold staining for RNAs and Western blotting for La. (E) Permeabilization buffer contains La-bound pre-tRNAs. Northern blotting for $tRNA^{Tyr\text{-}GTA}$ and $tRNA^{iMet\text{-}CAT}$ are shown.

Supplementary Figure S10

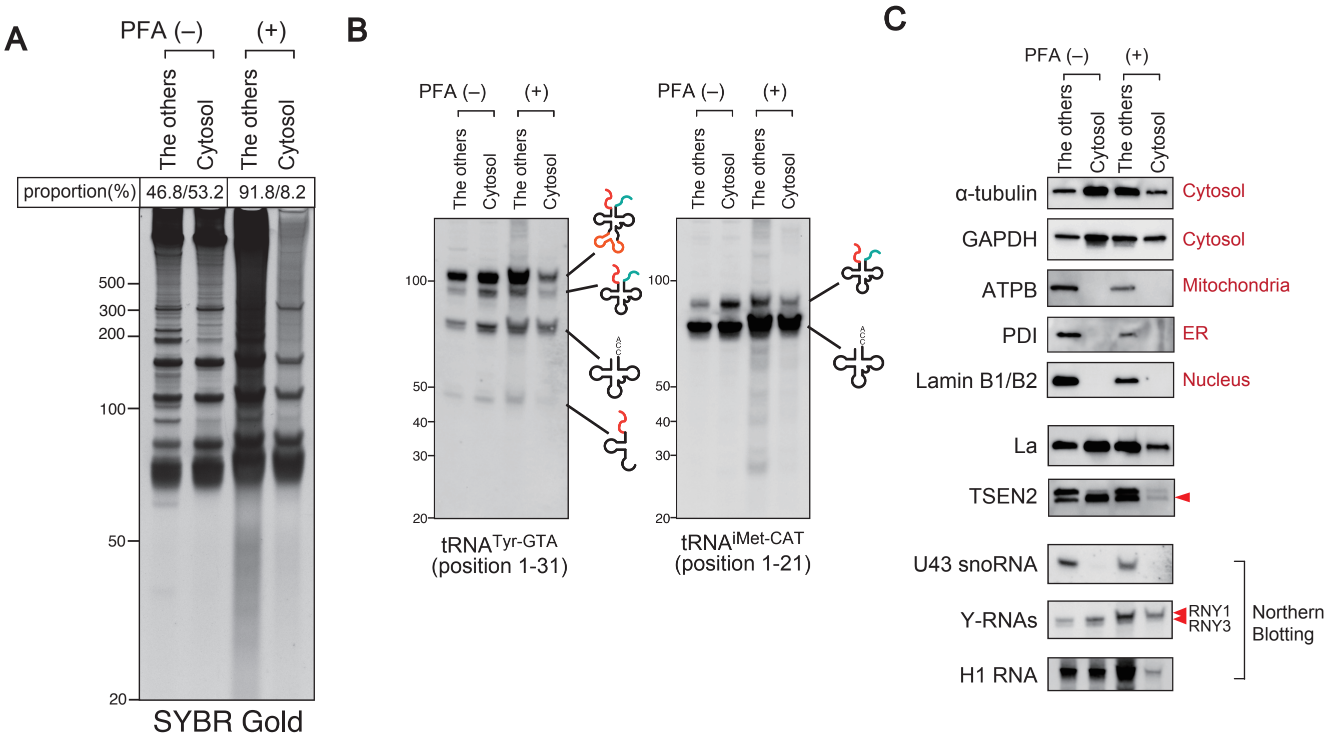


**Supplementary Figure S10.** Cytosol contains pre-tRNAs. The cytosolic fraction was obtained by digitonin-based subcellular fractionation combined with pre-treatment of the cells with PFA to minimize the leakage of nuclear content during fractionation by inhibiting the active nuclear-cytoplasmic transport. (A) RNA content of each fraction. SYBR Gold staining and the proportion of each fraction are shown. (B) Northern blotting for $tRNA^{Tyr\text{-}GTA}$ and $tRNA^{iMet\text{-}CAT}$. Even under the condition where nuclear export was inhibited by PFA fixation, the cytosolic fraction contains substantial proportion of pre-tRNAs. (C) Western blotting and Northern blotting for various molecules including nuclear, cytoplasmic and organelle markers.

### Supplementary Figure S11

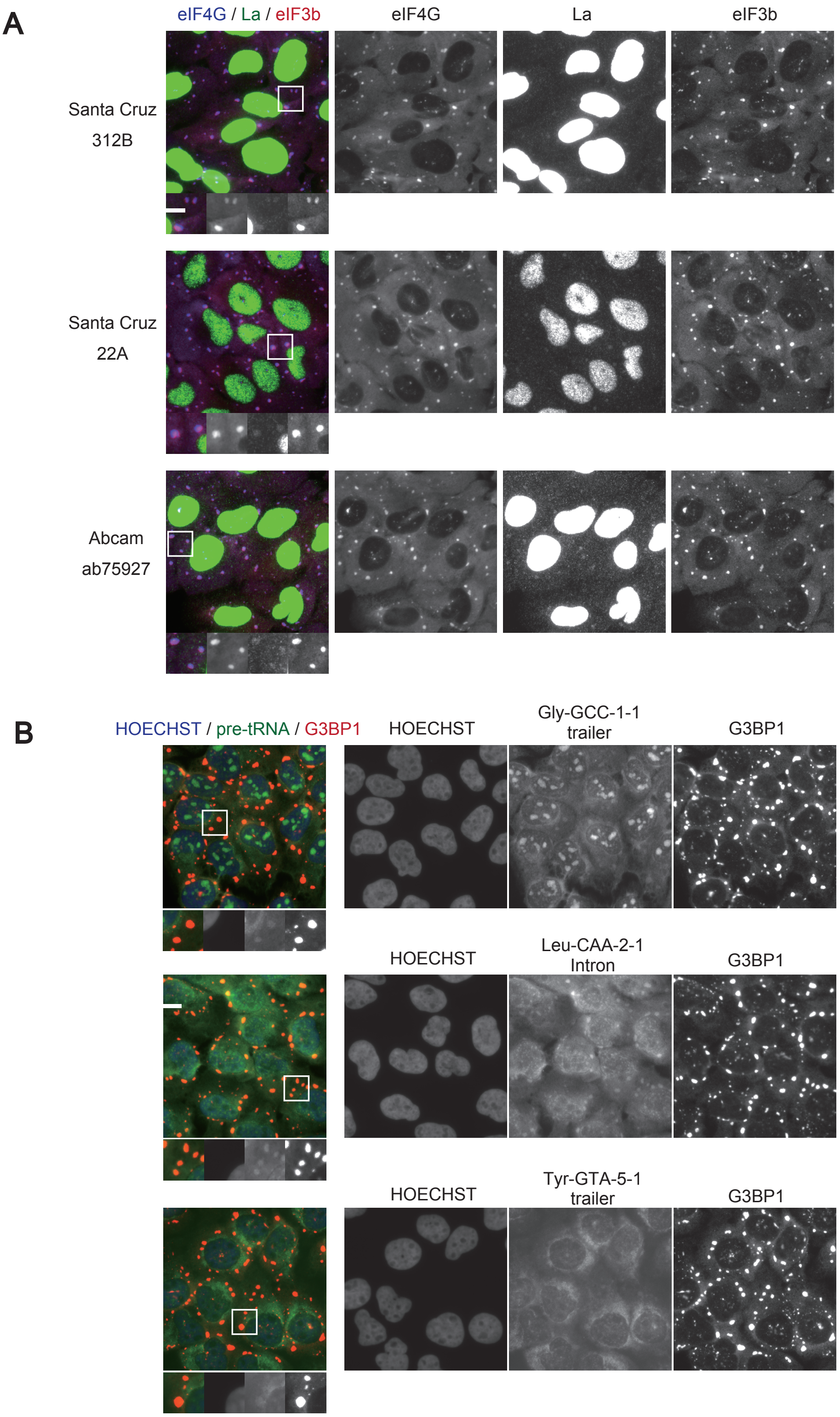


**Supplementary Figure S11.** La and pre-tRNAs are incorporated into stress granules (SGs).
(A) Immunofluorescence for La under stress condition. SGs are induced by incubation with 200 μM sodium arsenite for 1 hr. (B) pre-tRNAs are also incorporated into SGs. pre-tRNA FISH was performed for 3 kinds of pre-tRNAs.

### Supplementary Figure S12

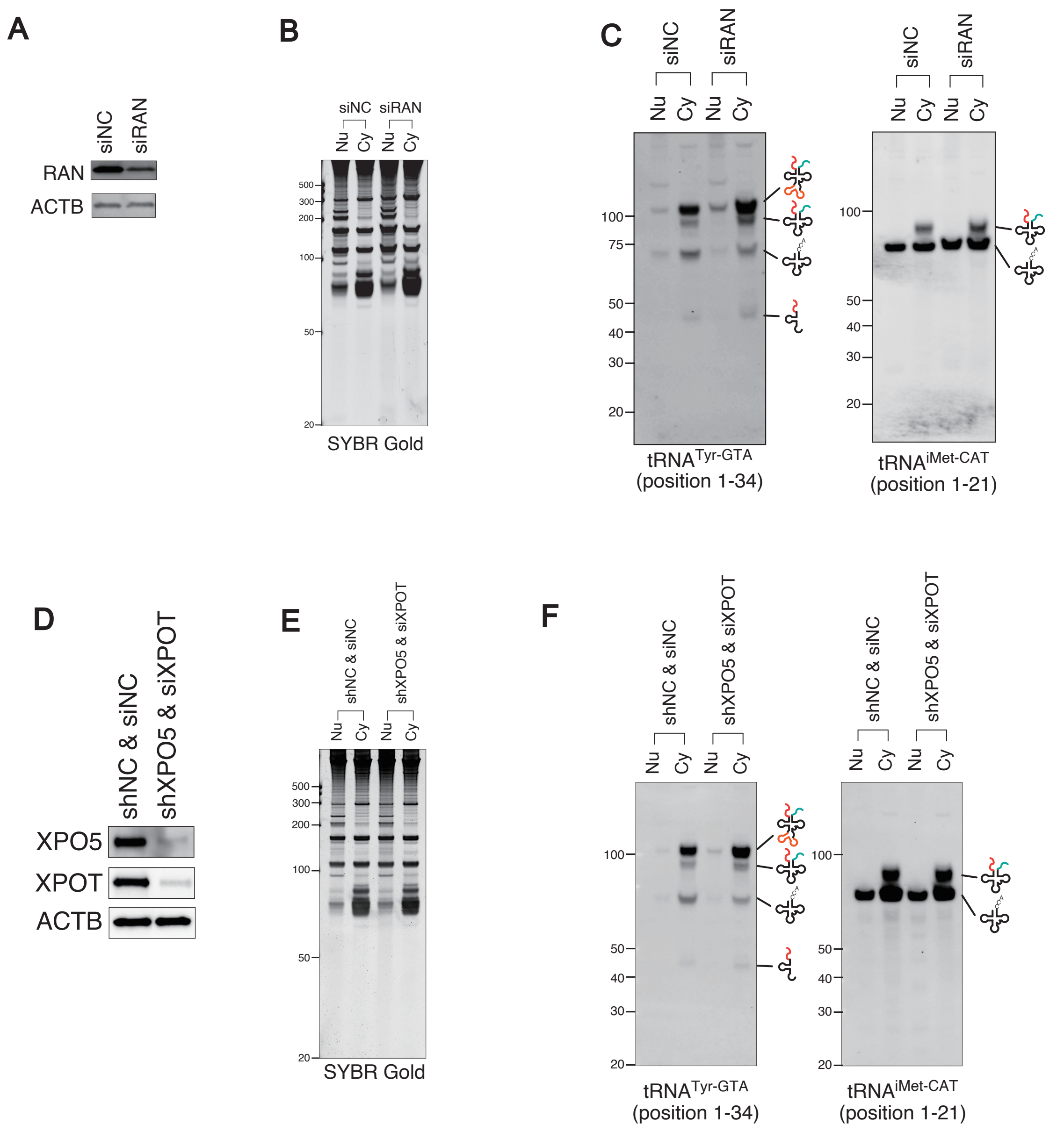


**Supplementary Figure S12.** Neither RAN knockdown nor knockdown of RNA-mediated tRNA exporters affects the localization of pre-tRNAs. (A-C) RAN knockdown does not affect the localization of pre-tRNAs. (A) Western blotting showing knockdown of RAN. (B) SYBR Gold staining of each fraction. (C) Northern blotting for $tRNA^{Tyr\text{-}GTA}$ and $tRNA^{iMet\text{-}CAT}$. Knockdown does not affect the localization of pre-tRNAs. (D-F) Double knockdown of XPO5 and XPOT does not affect the localization of pre-tRNAs. (D) Western blotting showing knockdown of XPO5 and XPOT. (E) SYBR Gold staining of each fraction. (F) Northern blotting for $tRNA^{Tyr\text{-}GTA}$ and $tRNA^{iMet\text{-}CAT}$. Knockdown does not affect the localization of pre-tRNAs.

Supplementary Figure S13

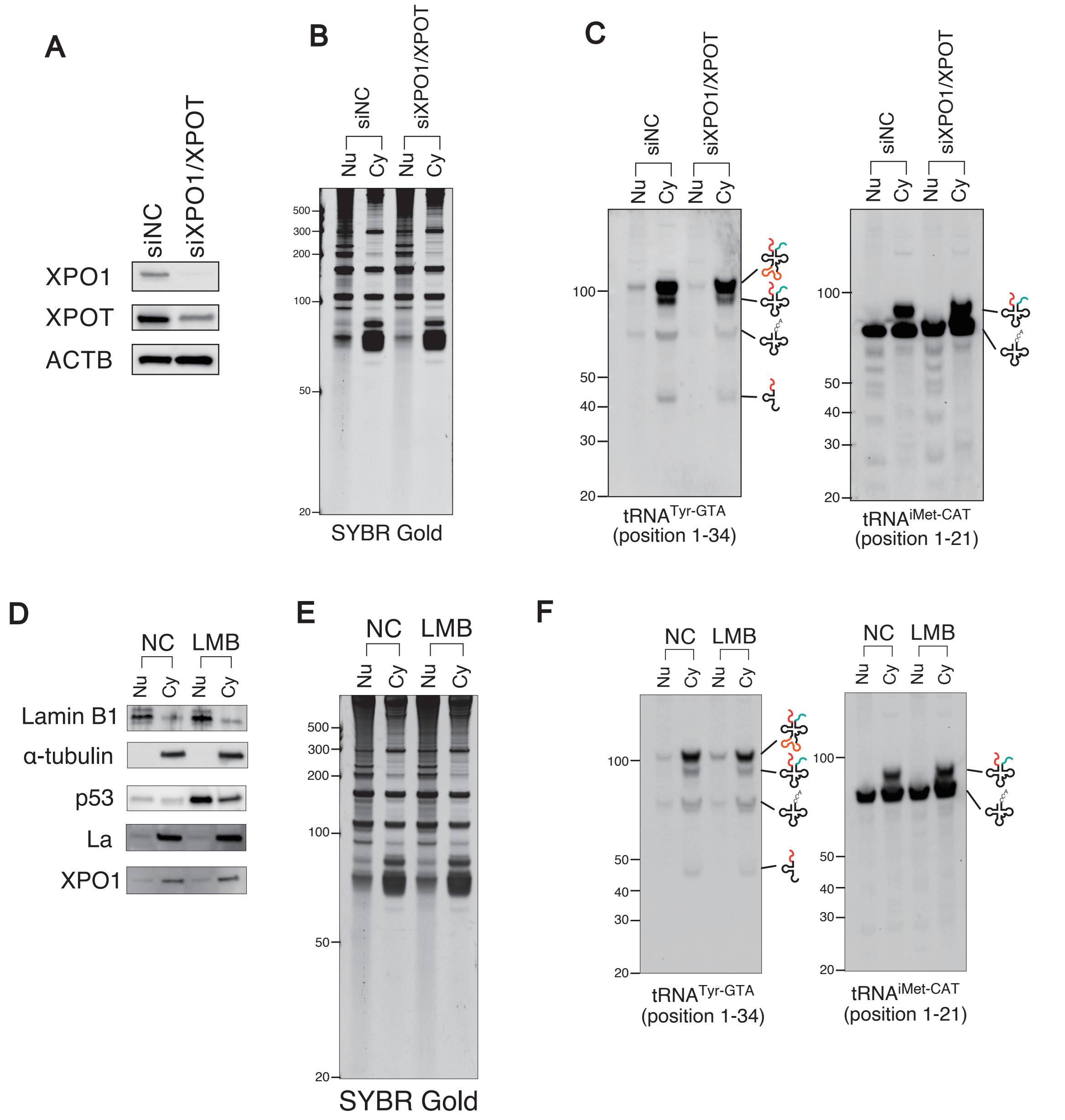


**Supplementary Figure S13.** Inhibition of XPO1 does not affect the localization of pre-tRNAs. (A-C) Double knockdown of XPO1 and XPOT does not affect the localization of pre-tRNAs. (A) Western blotting showing knockdown of XPO1 and XPOT. (B) SYBR Gold staining of each fraction. (C) Northern blotting for $tRNA^{Tyr\text{-}GTA}$ and $tRNA^{iMet\text{-}CAT}$. Knockdown does not affect the localization of pre-tRNAs. (D-F) Leptomycin B-mediated XPO1 inhibition does not affect the localization of pre-tRNAs. U2OS cells were treated with 10 ng/ml Leptomycin B (LMB) for 8 h. (D) Western blotting, (E) SYBR Gold staining and (F) Northern blotting for $tRNA^{Tyr\text{-}GTA}$ and $tRNA^{iMet\text{-}CAT}$. Note that LMB treatment induced the accumulation of p53 in the nuclear fraction, suggesting that LMB inhibited nuclear-cytoplasmic export. Under the condition, the localization of pre-tRNAs were not affected by LMB treatment.

### Supplementary Figure S14

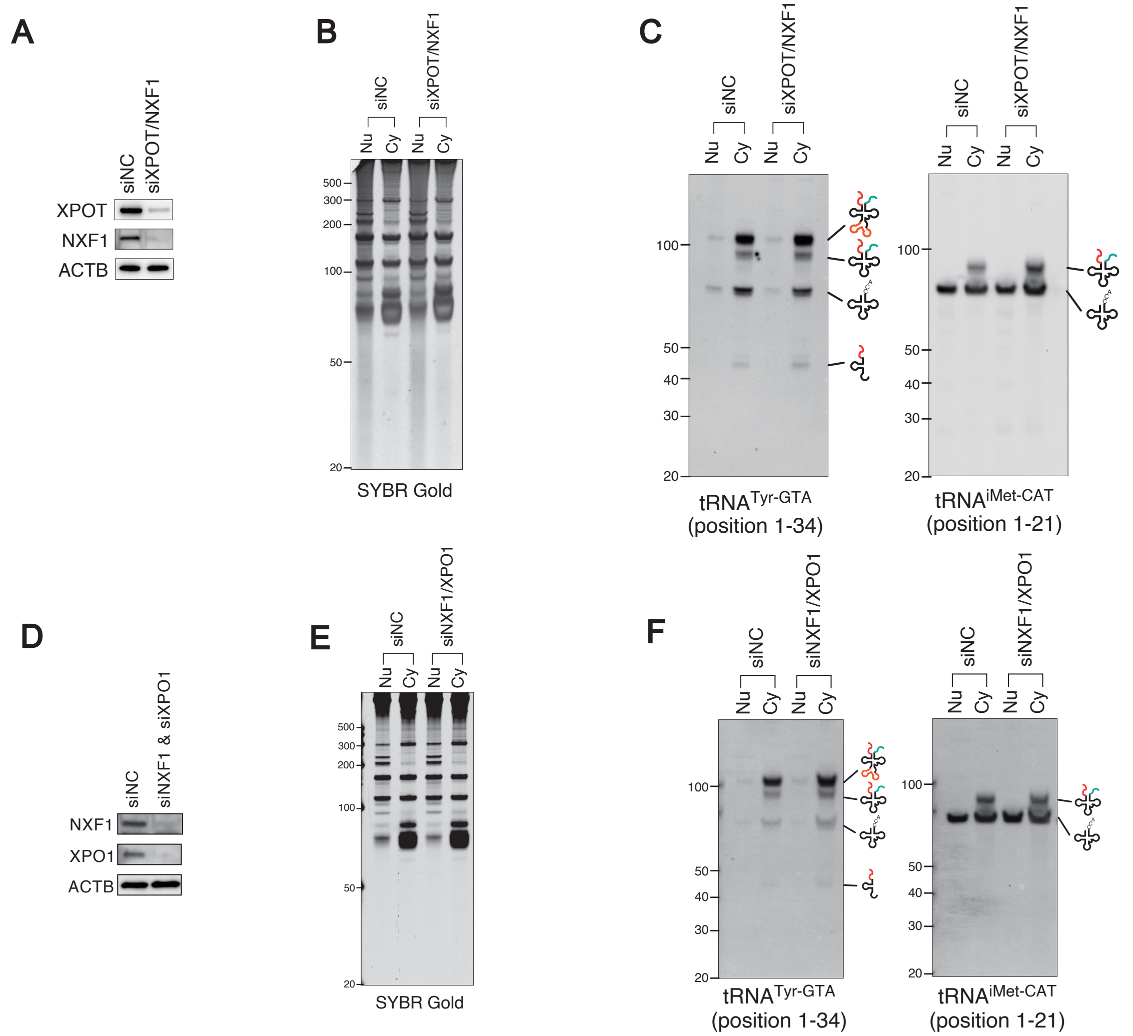


**Supplementary Figure S14.** NXF1 knockdown does not affect the localization of pre-tRNAs. (A-C) Double knockdown of NXF1 and XPOT does not affect the localization of pre-tRNAs. (A) Western blotting showing knockdown of NXF1 and XPOT. (B) SYBR Gold staining of each fraction. (f) Northern blotting for tRNA$^{Tyr\text{-}GTA}$ and tRNA$^{iMet\text{-}CAT}$. (D-F) Double knockdown of NXF1 and XPO1 does not affect the localization of pre-tRNAs. (D) Western blotting showing knockdown of NXF1 and XPO1. (E) SYBR Gold staining of each fraction. (F) Northern blotting for tRNA$^{Tyr\text{-}GTA}$ and tRNA$^{iMet\text{-}CAT}$.

### Supplementary Figure S15

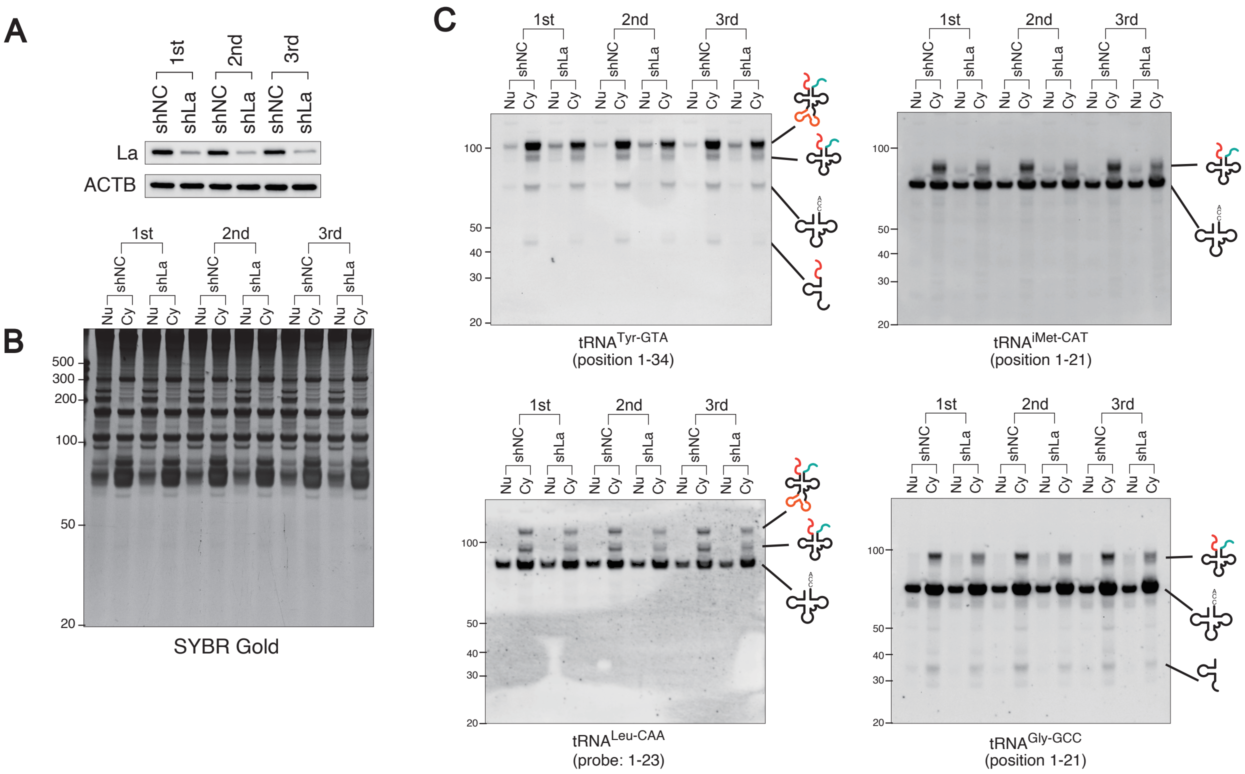


**Supplementary Figure S15.** Whole image of Figure 5. (A) Western blotting for La. (B) SYBR Gold staining of each fraction. (C) Northern blotting for $tRNA^{Tyr\text{-}GTA}$ and $tRNA^{iMet\text{-}CAT}$. Northern blotting for other tRNAs ($tRNA^{Leu\text{-}CAA}$ and $tRNA^{Gly\text{-}GCC}$) are also shown.

### Supplementary Table S1

| | | sequences |
|---|---|---|
| shNC (negative control) | top strand | ccggGCATTCACTTGGATAGTAActcgagTTACTATCCAAGTGAATGCttttt |
| | bottom strand | aattaaaaaGCATTCACTTGGATAGTAActcgagTTACTATCCAAGTGAATGC |
| shTSEN2 #1 | top strand | ccggGCTAAATGGAAAGATATGAAGctcgagCTTCATATCTTTCCATTTAGCttttt |
| | bottom strand | aattaaaaaGCTAAATGGAAAGATATGAAGctcgagCTTCATATCTTTCCATTTAGC |
| shTSEN2 #2 | top strand | ccggGATGTTTAAGTATTTACTATGctcgagCATAGTAAATACTTAAACATCttttt |
| | bottom strand | aattaaaaaGATGTTTAAGTATTTACTATGctcgagCATAGTAAATACTTAAACATC |
| shTSEN2 #3 | top strand | ccggGTTGTAATCGTCCATTAATTCctcgagGAATTAATGGACGATTACAACttttt |
| | bottom strand | aattaaaaaGTTGTAATCGTCCATTAATTCctcgagGAATTAATGGACGATTACAAC |
| shLa | top strand | ccggGCAAATGAGATTTCTTTGAATctcgagATTCAAAGAAATCTCATTTGCttttt |
| | bottom strand | aattaaaaaGCAAATGAGATTTCTTTGAATctcgagATTCAAAGAAATCTCATTTGC |
| shXPO5 | top strand | ccggGTAGAAACACCATCAAACTTTctcgagAAAGTTTGATGGTGTTTCTACttttt |
| | bottom strand | aattaaaaaGTAGAAACACCATCAAACTTTctcgagAAAGTTTGATGGTGTTTCTAC |

**Supplementary Table S1.** The sequences of DNA-oligos inserted to pLKO1 vector for shRNA-mediated knockdown.

Supplementary Table S2

| Probes | Sequences |
|---|---|
| U43 | 5'-AGCACACAGTTTCTGTCCGCCCG-3' |
| snR39B | 5'-GGTCAGTCCCGAAAGATGATTGCC-3' |
| HBII-202 | 5'-GGCCGTACAGCGATTCCGGAGAA-3' |
| RNY1 | 5'-GGAGTTCGATCTGTAACTGACTGTG-3' |
| RNY3 | 5'-ACACCACTGCACTCGGACCAGCC-3' |
| tRNA-Tyr-GTA(1-34) | 5'-CAGTCCTCCGCTCTACCAACTGAGCTATCGAAGG-3' |
| tRNA-Tyr-GTA(50-73) | 5'-TCCTTCGAGCCGGAATCGAACCA-3' |
| tRNA-Tyr-GTA-2-1 5'-Leader | 5'-TGAGCTATCGAAGGCTCCGCT-3' |
| tRNA-Tyr-GTA-2-1 Intron | 5'-TGCCACGCCCTATCCACTACA-3' |
| tRNA-Tyr-GTA-5-1 3'-Trailer | 5'-AAAACCGCACTTGTCTCCTTCGA-3' |
| tRNA-Arg-TCT(1-23) | 5'-GCTATCCATTGCGCCACAGAGCC-3' |
| tRNA-Arg-TCT(51-73) | 5'-CGACTCTGGTGGGACTCGAACCC-3' |
| tRNA-Leu-CAA(1-23) | 5'-TTAGACCACTCGGCCATCCTGAC-3' |
| tRNA-Leu-CAA(50-73) | 5'-TGTCAGAAGTGGGATTCGAACCCA-3' |
| tRNA-Leu-CAA-2-1 Intron | 5'-GACCCGAACACAGGAAGCAGTAAG-3' |
| tRNA-iMet-CAT(1-21) | 5'-CTTCCGCTGCGCCACTCTGCT-3' |
| tRNA-Gly-GCC(1-21) | 5'-CTACCACTGAACCACCCATGC-3' |
| tRNA-Gly-GCC-1-1 3'-Trailer | 5'-AAATGGGAGGGCGTGCTGCAT-3' |
| H1 RNA (134-153) | 5'-CCCGTTCTCTGGGAACTCAC-3' |
| H1 RNA (160-183) | 5'-TCTCCTGCCCAGTCTGACCTCGCG-3' |
| H1 RNA (240-261) | 5'-GCTGGCCGTGAGTCTGTTCCAA-3' |
| H1 RNA (318-340) | 5'-AATGGGCGGAGGAGAGTAGTCTG-3' |

**Supplementary Table S2.** The sequences of DNA-oligo probes for Northern blotting.

Supplementary Table S3

| Antibodies | Source | Identifier |
|---|---|---|
| Rabbit anti-Lamin B1 | Santa Cruz | sc-20682 |
| Mouse anti-Lamin B1/B2 | ThermoFischer | MA1-90041 |
| Mouse anti-α-tubulin | Proteintech | 66031-1-Ig |
| Mouse anti-ACTB | Proteintech | 66009-1-Ig |
| Mouse anti-GAPDH | Santa Cruz | sc-47724 |
| Mouse anti-ATPB | MitoSciences | MS503 |
| Rabbit anti-PDI | Cell Signaling | #2446 |
| Rabbit anti-TSEN2 | Novus | NBP1-81140 |
| Rabbit anti-TSEN2 | Proteintech | 13103-2-AP |
| Rabbit anti-TSEN2 | GeneTex | GTX32940 |
| Rabbit anti-TSEN2 | Sigma-Aldrich | HPA027120 |
| Mouse anti-TSEN34 | ORIGENE | TA810146 |
| Mouse anti-TSEN15 | Santa Cruz | sc-374085 |
| Mouse anti-TSEN54 | Santa Cruz | sc-374488 |
| Rabbit anti-CLP1(HEAB) | Abcam | ab133669 |
| Mouse anti-ARCH | ThermoFischer | MA5-24536 |
| Rabbit anti-DDX1 | Santa Cruz | sc-134752 |
| Goat anti-RTCB | LSBio | LS-C139807 |
| Mouse anti-RTCB(HSPC117/FAAP) | Santa Cruz | sc-393966 |
| Rabbit anti-POP4 | Proteintech | 15273-1-AP |
| Rabbit anti-Rpp21 | Proteintech | 16377-1-AP |
| Mouse anti-RNase Z1 | Santa Cruz | sc-390029 |
| Rabbit anti-TRNT1 | Novus | NBP1-86589 |
| Mouse anti-La/SSB (312B) | Santa Cruz | sc-80656 |
| Mouse anti-La/SSB (22A) | Santa Cruz | sc-80655 |
| Mouse anti-SSB | Abcam | ab75927 |
| Rabbit anti-SSB | Novus | NBP1-33549 |
| Mouse anti-La/SSB | Santa Cruz | sc-80656 |
| Mouse anti-XPOT | Santa Cruz | sc-514591 |
| Mouse anti-XPO5 | Santa Cruz | sc-271036 |
| Mouse anti-XPO1 | Santa Cruz | sc-74454 |
| Rabbit ant-p53 | Leica Biosystems | NCL-p53-CM1 |
| Mouse anti-NXF1(TAP) | Santa Cruz | sc-32319 |
| Mouse anti-RAN | Santa Cruz | sc-271376 |
| Mouse IgG1κ | BD Pharmingen | 557273 |

**Supplementary Table S3.** The list of antibodies used in this study.

Supplementary Table S4

| Probes | Sequences |
|---|---|
| tRNA-Leu-CAA-2-1 Intron | 5'-CTTACTGCTTCCTGTGTTCGGGTC-Bio-3' |
| tRNA-Gly-GCC-1-1 3'-trailer | 5'-AAATGGGAGGGCGTGCTGCAT-Bio-3' |
| tRNA-Tyr-GTA-5-1 3'-trailer | 5'-AAAACCGCACTTGTCTCCTTCGA-Bio-3' |
| H1 RNA (134-153) | 5'-CCCGTTCTCTGGGAACTCAC-Bio-3' |
| H1 RNA (160-183) | 5'-TCTCCTGCCCAGTCTGACCTCGCG-Bio-3' |
| H1 RNA (240-261) | 5'-GCTGGCCGTGAGTCTGTTCCAA-Bio-3' |

**Supplementary Table S4.** The sequences of DNA-oligo probes for fluorescent *in situ* hybridization (FISH) used in this study.
